# Calcium signaling in the photodamaged skin

**DOI:** 10.1101/2021.05.14.443726

**Authors:** Viola Donati, Chiara Peres, Chiara Nardin, Ferdinando Scavizzi, Marcello Raspa, Catalin D. Ciubotaru, Mario Bortolozzi, Fabio Mammano

**Affiliations:** Department of Physics and Astronomy “G. Galilei”, University of Padova, 35131 Padova, Italy; CNR Institute of Biochemistry and Cell Biology, 00015 Monterotondo, Rome, Italy; CNR IOM Materials Foundry, 34149 Basovizza (TS), Italy; Veneto Institute of Molecular Medicine, Viale Giuseppe Orus 2, 35131 Padova, Italy

**Keywords:** Skin, laser-induced intra-epidermal photodamage, wound healing, purinergic signaling, connexins, pannexins

## Abstract

**BACKGROUND:** The mammalian skin, the body’s largest single organ, is a highly organized tissue that forms an essential barrier against dehydration, pathogens, light and mechanical injury. Damage triggers perturbations of the cytosolic free Ca^2+^ concentration ([Ca^2+^]_c_) that spread from cell to cell (known as intercellular Ca^2+^ waves) in different epithelia, including epidermis. Ca^2+^ waves are considered a fundamental mechanism for coordinating multicellular responses, however the mechanisms underlying their propagation in the damaged epidermis are incompletely understood.

**AIM OF THE PROJECT:** To dissect the molecular components contributing to Ca^2+^ wave propagation in murine model of epidermal photodamage.

**METHODS:** To trigger Ca^2+^ waves, we used intense and focused pulsed laser radiation and targeted a single keratinocyte of the epidermal basal layer in the earlobe skin of live anesthetized mice. To track photodamage-evoked Ca^2+^ waves, we performed intravital multiphoton microscopy in transgenic mice with ubiquitous expression of the sensitive and selective Ca^2+^ biosensor GCaMP6s. To dissect the molecular components contributing to Ca^2+^ wave propagation, we performed *in vivo* pharmacological interference experiments by intradermal microinjection of different drugs.

**EXPERIMENTAL RESULTS:** The major effects of drugs that interfere with degradation of extracellular ATP or P2 purinoceptors suggest that Ca^2+^ waves in the photodamaged epidermis are primarily due to release of ATP from the target cell, whose plasma membrane integrity was compromised by laser irradiation. The limited effect of the Connexin 43 (Cx43) selective inhibitor TAT-Gap19 suggests ATP-dependent ATP release though connexin hemichannels (HCs) plays a minor role, affecting Ca^2+^ wave propagation only at larger distances, where the concentration of ATP released from the photodamaged cell was reduced by the combined effect of passive diffusion and hydrolysis due to the action of ectonucleotidases. The ineffectiveness of probenecid suggests pannexin channels have no role. As GCaMP6s signals in bystander keratinocytes were augmented by exposure to the Ca^2+^ chelator EGTA in the extracellular medium, the corresponding transient increments of the [Ca^2+^]_c_ should be ascribed primarily to Ca^2+^ release from the ER, downstream of ATP binding to P2Y purinoceptors, with Ca^2+^ entry through plasma membrane channels playing a comparatively negligible role. The effect of thapsigargin (a well-known inhibitor of SERCA pumps) and carbenoxolone (a recently recognized inhibitor of Ca^2+^ release through IP_3_ receptors) support this conclusion.

**CONCLUSIONS:** The one presented here is an experimental model for accidental skin injury that may also shed light on the widespread medical practice of laser skin resurfacing, used to treat a range of pathologies from photodamage and acne scars to hidradenitis suppurativa and posttraumatic scarring from basal cell carcinoma excision. The results of our experiments support the notion that Ca^2+^ waves reflect chiefly the sequential activation of bystander keratinocytes by the ATP released through the compromised plasma membrane of the cell hit by laser radiation. We attributed the observed increments of the [Ca^2+^]_c_ chiefly to signal transduction through purinergic P2Y receptors. Several studies have highlighted fundamental roles of P2Y receptors during inflammatory and infectious diseases, and the initial phase of wound healing involves acute inflammation. In addition, hyaluronan is a major component of the extracellular matrix and its synthesis is rapidly upregulated after tissue wounding via P2Y receptor activation. It is tempting to speculate that response coordination after injury in the epidermis occurs via propagation of the ATP-dependent intercellular Ca^2+^ waves described in this work.

## INTRODUCTION

The mammalian skin, the body’s largest single organ, is a highly organized tissue that comprises three different substructures, the epidermis, the dermis and the hypodermis, and forms an essential barrier against dehydration, pathogens, light and mechanical injury [1]. The epidermis, the body’s interface to the environment, is a stratified epithelium largely composed of keratinocytes (95%), melanocytes (that donate pigment to the keratinocytes), Langerhans’ cells, which have immunological functions, and Merkel cells, also known as Merkel-Ranvier cells or tactile epithelial cells [2]. It is well established that epidermis’ continuous renewal is sustained by stem cells contained in the *stratum basale* [3-7]. In the epidermis, the differentiating keratinocytes are gradually displaced in the outward direction, from *stratum basale* through *stratum spinosum* to *stratum corneum*, where corneocytes are continually shed from the skin surface.

A gradient of Ca^2+^ across the epidermis [8] is key for keratinocyte differentiation and formation of the epidermal permeability barrier [9, 10], maintenance of which relies on a delicate balance between proliferation and differentiation [2, 11], two Ca^2+^-dependent cellular processes [12]. Defects of epidermal Ca^2+^ homeostasis cause skin pathologies, such as Darier’s disease for which *ATP2A2* has been identified as the defective gene indicating that the sarcoplasmic/endoplasmic reticulum Ca^2+^ ATPase (SERCA)2 plays a key role [13].

ATP, which is present at mM concentration in the cell cytosol whereas its normal concentration in the extracellular environment is in low nM range, is released by most cells as an extracellular signaling molecule [14, 15]. ATP release was detected in cultured human neonatal keratinocytes exposed to air [16] as well as from injured mouse epidermal keratinocytes [17], HaCaT cells [18] (a spontaneously transformed aneuploid immortal keratinocyte cell line from adult human skin, with high capacity to differentiate and proliferate in vitro [19]) and normal human epidermal keratinocytes (NHEKs) co-cultured with mouse dorsal root ganglion (DRG) neurons [20]. Extracellular ATP can affect the cytosolic free Ca^2+^ concentration ([Ca^2+^]_C_) by activating Ca^2+^-permeable P2X receptors [21] and/or by promoting Ca^2+^-release from the endoplasmic reticulum (ER) via G-protein coupled P2Y receptors [22, 23] on the surface of keratinocytes and other epidermal cells types [24]. Therefore, ATP release and purinergic signaling have the potential to interfere with epidermal homeostasis and have been implicated in a host of processes, including pain, inflammation and wound healing [24].

In different epithelia, including epidermis, damage triggers perturbations of the [Ca^2+^]_c_ that spread from cell to cell (known as intercellular Ca^2+^ waves) [25-30]. Ca^2+^ waves are considered a fundamental mechanism for coordinating multicellular responses [31], however the mechanisms underlying Ca^2+^ wave propagation and their significance in the damaged epidermis are incompletely understood. Here, we studied intercellular Ca^2+^ waves evoked by selectively photodamaging a single keratinocyte of the epidermal basal layer of the mouse earlobe skin. Our experimental model is related not only to accidental injury, but also to the medical practice of skin resurfacing/rejuvenation based on intra-epidermal focal laser-induced photodamage [32-35]. To visualize Ca^2+^ waves, we used intravital multiphoton microscopy in anesthetized transgenic mice expressing the sensitive and selective cytosolic Ca^2+^ biosensor GCaMP6s [36]. To dissect the molecular components contributing to Ca^2+^ wave propagation, we performed *in vivo* pharmacological interference experiments by intradermal microinjection of different drugs.

## MATERIALS AND METHODS

### Study design

To estimate the minimum sample size of each experimental group for animal experiments, we set a probability *α* = 5% for the type I error in the ANOVA test. Then, fixing β = 4α = 20% to obtain a test power of 1 − β = 80%, we computed the number of samples *n* using the formula:

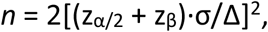

with z_α/2_ = 1.96 and z_β_ = 0.8416. We assumed σ = 0.35 (35%) as estimation for variability of data (σ^2^ variance of data distribution) and Δ = 0.75 as expected effect size (minimum normalized difference between averages that has biological significance). The experiment was repeated until a minimum number of samples was collected for each experimental group. We stopped data collection when at least *n* = 3 photodamage experiments were performed in at least *m* = 1 mice for each experimental condition and pooled the results. The investigator was not blinded during administration of treatments or result assessment. No samples were excluded from analyses.

### Animals

Mice were bred and genotyped at the National Research Council-Institute of Biochemistry and Cell Biology (CNR-IBBC), Infrafrontier/ESFRI-European Mouse Mutant Archive (EMMA), Specific Pathogen-Free (SPF) barrier unit (Monterotondo Scalo, Rome). All the experimental procedures were agreed upon, reviewed and approved by local animals welfare oversight bodies and were performed with the approval and direct supervision of the CNR-IBBC/Infrafrontier—Animal Welfare and Ethical Review Body (AWERB), in accordance with general guidelines regarding animal experimentation, approved by the Italian Ministry of Health, in compliance with the Legislative Decree 26/2014 (ref. Project license 603/2018-PR), transposing the 2010/63/EU Directive on protection of animals used in research). In addition, all animal experimentation was conducted in adherence to the NIH Guide for the Care and Use of Laboratory Animals and recommendations from both ARRIVE and PREPARE guidelines [37, 38]. Mice were housed in individually ventilated caging systems (Tecniplast, Gazzada, Italy) at a temperature (T) of 21 ± 2 °C, relative humidity (RH) of 55 ± 15% with 50–70 air changes per hour (ACH) and under controlled (12 : 12 hour) light–dark cycles (7 am–7 pm). Mice had ad libitum access to water and a standard rodent diet (Emma 23, Mucedola, Settimo Milanese, Italy).

Adult transgenic mice, both male and female, aged between 5 and 40 weeks and ubiquitously expressing the Ca^2+^ biosensor GCaMP6s [36] were used for in vivo Ca^2+^ imaging experiments. These mice were generated in the animal facility of the laboratory by crossing the Jakson Laboratory strain #024106 (STOCK-Gt(ROSA)26Sor^*tm*96(*CAG*−*GCaMP6s*)*Hze*^ /J) with the European Mouse Mutant Archive (EM) Cre-deleter strain EM:01135 (B6.C-Tg(CMVcre) ^1*Cgn/CgnCnrm*^). Double mutant mice were identified amplifying tail genomic DNA by means of PCR. The presence of the CAG-GCamp6 insertion in the Rosa26 locus and the wild type allele were detected using the primers:

5′-ACG-AGT-CGG-ATC-TCC-CTT-TG-3′, 5′-AAG-GGA-GCT-GCA-GTG-GAG-TA-3′

and

5′-CCG-AAA-ATC-TGT-GGG-AAG-TC-3′.

The CRE-deleter transgene was detected using the primer pair

5′-CGA-GTG-ATG-AGG-TTC-GCA-AG -3′

and

5′-TGA-GTG-AAC-GAA-CCT-GGT-CG -3′.

PCR products were run on 2% agarose gels, visualized with ethidium bromide and photographed. Expected band sizes were 450 bp for the CAG-GCamp6 and 297 bp for the wt allele and 390bp for the Cre transgene.

### Multiphoton microscopy

#### System description

To carry out this experimental work, we used a custom-made multiphoton system (**Figure 1**) based on a Bergamo II architecture (Thorlabs Imaging System, Sterling, VI, USA), as previously described [29]. The system was equipped with two scanning heads, one with resonant-galvo (RG) mirrors and the other with galvo-galvo (GG) mirrors, and was coupled to a mode-locked titanium-sapphire (Ti:Sa) fs pulsed laser (Chameleon Vision II Laser, Coherent, Inc., Santa Clara, CA, USA) (**Figure 2**). The RG scanner was used for imaging, whereas the GG scanner was used to focally photodamage a pre-defined spot in the field of view (FOV) by focusing the collimated laser beam onto the sample through a 25× water-immersion objective (XLPLN25XWMP2, NA 1.05, Olympus Corporation, Tokyo, Japan; the same objective was also used for imaging).

**Figure 1:**
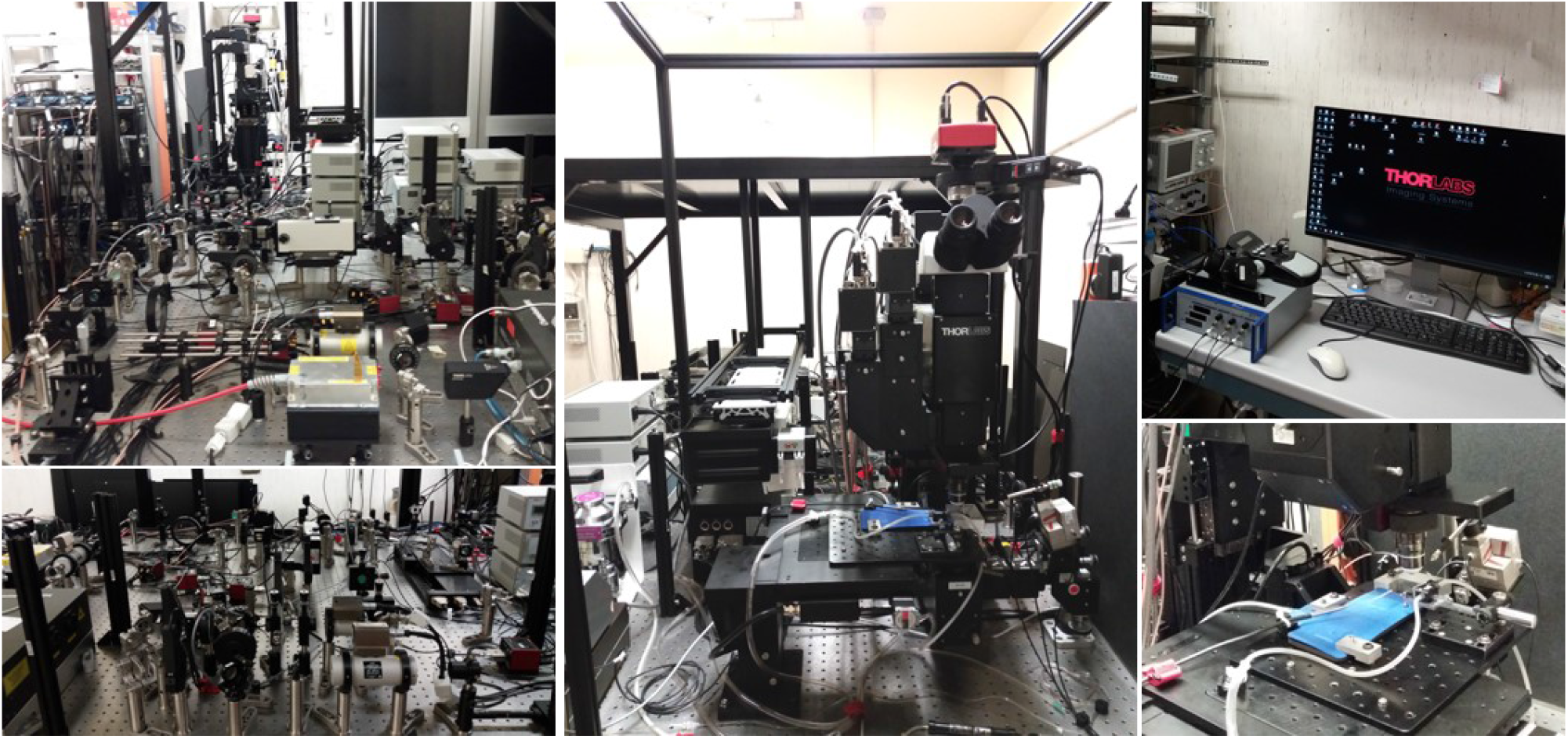
Photographs of the multiphoton-STED intravital microscopy facility at CNR, Rome.

**Figure 2:**
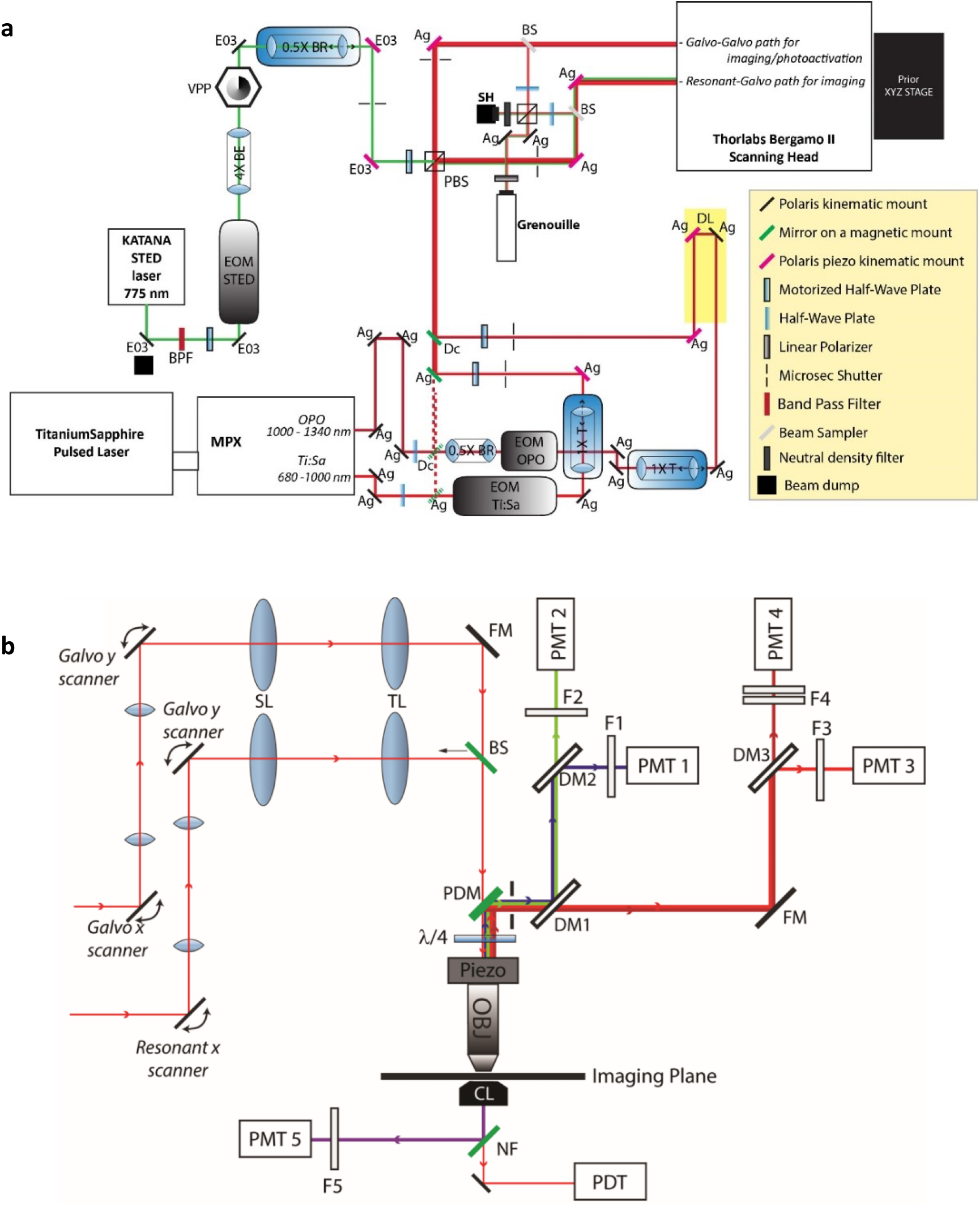
Simplified optical scheme of the optical paths and multiphoton-STED microscope. (**a**) Optical bench optics used to deliver 3 laser lines to the two scanning heads of the multiphoton microscope. VPP, vortex phase plate; SH, Shack-Hartmann wavefront sensor; Grenouille, beam profiler; Ag, silver-coated mirrors; E03, dielectric mirrors; PBS, polarizing beam splitter; EOM, electro-optical modulators; BS, beam sampler; BE, beam expander; BR, beam reducer; T, variable telescope used to control beam divergence. DL, optical delay line, used to synchronize PUMP and OPO pulses. (**b**) Multiphoton microscope. Red lines entering from the left represent titanium-sapphire (Ti:Sa) laser beams impinging on the pair of scanning systems. SL: Scan Lens; TL: Tube Lens; FM: Full Mirror; BS: 50/50 Beam splitter; PDM: Primary Dichroic Mirror; λ/4: Quarter wave plate; Piezo: Piezo Objective Scanner; Obj: Objective lens; DM1: 565 nm long pass filter (T565lpxr); DM2: 495 nm long pass filter (T495lpxru); DM3: 652 long pass filter (FF652-Di01-25×36); F1: 460/50 nm band pass filter (ET460/50m-2p); F2: 525/40 nm band pass filter (FF02-525/40-25); F3: 612/69 band pass filter (FF01-612/69-25); F4: combination of 647 nm long pass filter (BLP01-647R-25) and 770 nm short pass filter (FF01-770/SP-25); NF: Notch Filter; PMT: photomultiplier tube; PDT: photodiode tube.

Multiphoton excitation of GCaMP6s was performed at 920 nm, whereas its emission signal was filtered by a single band-pass filter (Cat. No. FF02-525/40-25, Semrock/IDEX, Rochester, NY, USA) placed in front of a non-descanned GaAsP detector (Cat. No. H7422–50, Hamamatsu Photonics K.K., Shizuoka, Japan. Electro-optical modulators (EOM) and mechanical ultra-fast shutters were used to control both photodamage and imaging light exposure using the ThorImage LS 3.1 software (Thorlabs). Parameters used for photodamage and image acquisition are listed in **Table 1**. Laser excitation intensity and frame averaging were adjusted to minimize photobleaching and phototoxicity, while achieving enough signal to noise ratio and temporal resolution.

**Table 1:** Photodamage and image acquisition parameters.

|  |  |
| --- | --- |
| Excitation and photodamage wavelength | 920 nm |
| Photodamage optical power in the focal plane | 118 mW |
| Photodamage duration | 500 ms |
| Imaging average power at focal optical plane | 25 mW |
| Frame size (pixels) | 512 × 512 |
| Pixel dwell time | 81.4 ns |
| Raw frame rate | 45.7 fps |
| Number of averaged frames | 9 |
| Effective frame rate | 2.5 fps |
| Resolution | 0.62 $\mu\text{m}/\text{pixel}$ |

#### Intravital multiphoton microscopy and drug delivery

Mice were anesthetized with an intraperitoneal (i.p.) injection of physiological solution containing 90 mg/kg ketamine and 0.5 mg/kg medetomidine. If the experiment duration was longer than 2 hours, half-dose of the anesthesia was reinjected to avoid mouse awaking. Mice were positioned on the stage of the multiphoton microscope equipped with a heated pad kept at 35°C for homeothermic control. Ca^2+^ signals were recorded from the basal keratinocytes of the mouse pinna (earlobe) skin, fixed under the objective with double-sided tape. For optical coupling, a drop of phosphate buffered saline (PBS) solution was placed between the skin and the 25× water-immersion objective lens. The PBS solution was composed of 10 mM Phosphate (as sodium phosphates), 2.68 mM KCl and 140 mM NaCl.

After positioning the animal under the microscope objective lens, we waited 10 minutes to allow intracellular Ca^2+^ (mobilized during earlobe manipulation) to return to baseline (as judged by low levels of GCaMP6s fluorescence). For each mouse, control photodamage experiments were repeated at least 3 times with similar results in different positions of the earlobe. Next, to block or enhance a specific pathway the ear surface was gently cleaned and dried and a 4 µl solution containing a selected drug (**Table 2**) diluted in a PBS was microinjected in the earlobe skin using a 10 µl NANOFIL syringe (World Precision Instruments Inc., Sarasota, FL, USA) equipped with a gauge 33 needle. The microinjected PBS solution also contained 1.78 µM of fluorescent Dextran Texas Red (MW=70,000; single-photon Excitation/Emission wavelength=595/615 nm; two-photon excitation wavelength=920) to confirm that the drug-containing fluid reached the imaged area. Thereafter, the same photodamage protocol used in the previous control recording was used to acquire image sequences in the presence of the selected drug. A dedicated series of experiments was performed to verify that photodamage-evoked Ca^2+^ waves were insensitive to the presence of the microinjected fluorescent Dextran Texas Red. Due to its large molecular weight (MW), Dextran diffused slowly through the tight layers of keratinocytes and accumulated mainly in the extracellular spaces below the *stratum basale*.

**Table2:**
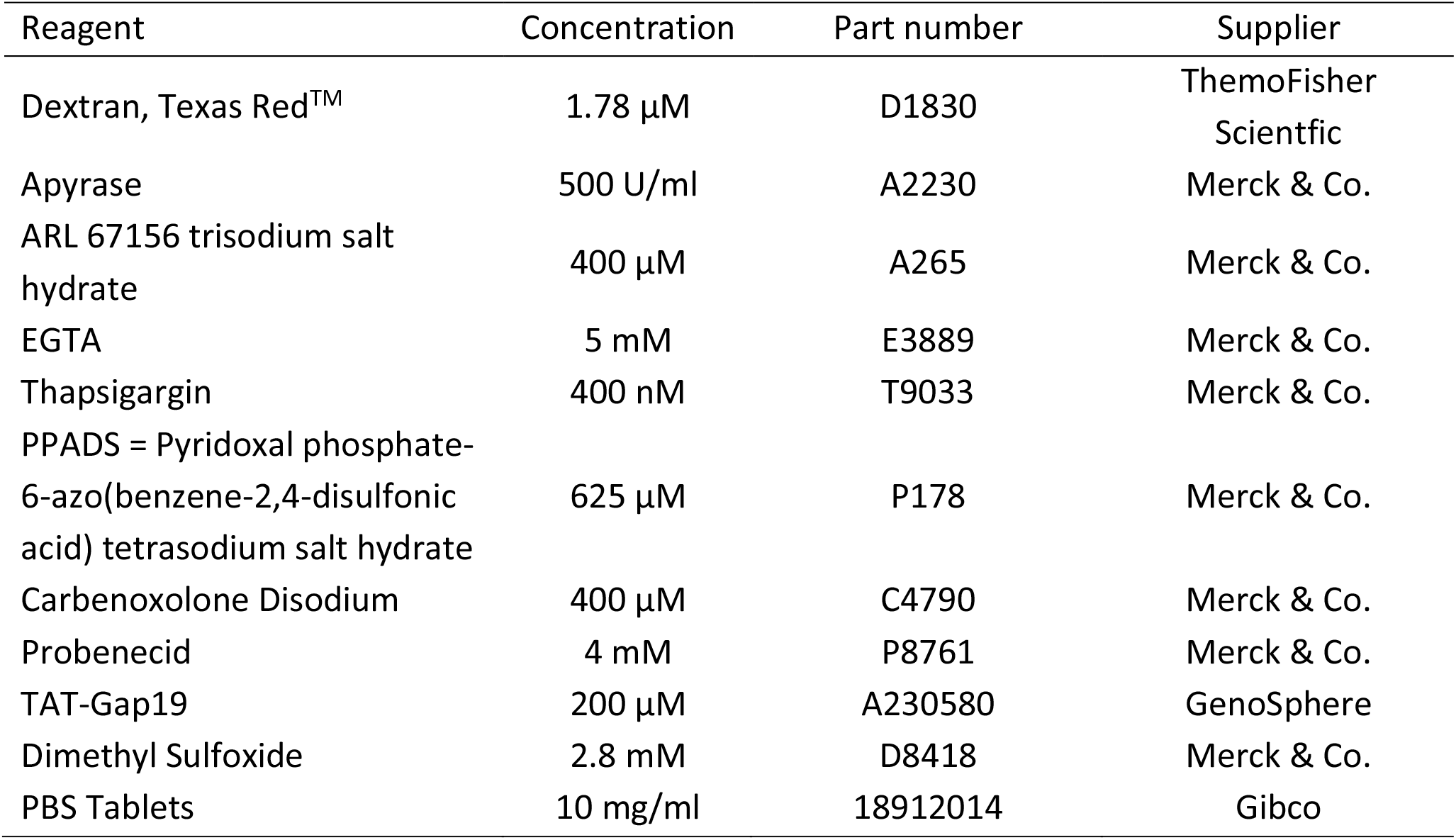
Drugs used to dissect components of Ca^2+^ wave propagation mechanisms.

| Reagent | Concentration | Part number | Supplier |
| --- | --- | --- | --- |
| Dextran, Texas Red™ | 1.78 µM | D1830 | ThermoFisher Scientific |
| Apyrase | 500 U/ml | A2230 | Merck & Co. |
| ARL 67156 trisodium salt hydrate | 400 µM | A265 | Merck & Co. |
| EGTA | 5 mM | E3889 | Merck & Co. |
| Thapsigargin | 400 nM | T9033 | Merck & Co. |
| PPADS = Pyridoxal phosphate-6-azo(benzene-2,4-disulfonic acid) tetrasodium salt hydrate | 625 µM | P178 | Merck & Co. |
| Carbenoxolone Disodium | 400 µM | C4790 | Merck & Co. |
| Probenecid | 4 mM | P8761 | Merck & Co. |
| TAT-Gap19 | 200 µM | A230580 | GenoSphere |
| Dimethyl Sulfoxide | 2.8 mM | D8418 | Merck & Co. |
| PBS Tablets | 10 mg/ml | 18912014 | Gibco |

In general, while imaging the skin, a given focal plane intercepts furrows and bulges, exposing a multitude of cell populations and structures. Cells corresponding to different epidermal layers can be distinguished morphologically: smaller polygonal cells are keratinocytes of the *stratum basale*, while increasingly larger cells correspond to the suprabasal layers. Images acquired with different PMTs of the multiphoton microscope are shown in **Figure 3** together with the corresponding composite images. Second-harmonic generation (SHG) images were acquired at 460/50 nm emission wavelength, allowing the imaging of collagen fibers (**Figure 3 a**,**e**). GCaMP6s fluorescence (and keratinocytes autofluorescence) contributed to the signals detected in the 525/40 nm emission channel (**Figure 3 b**,**f**). Images in **Figure 3 c**,**g** were acquired in the 612/90 nm emission channel before and after the microinjection of fluorescent Dextran Texas Red™.

**Figure 3:**
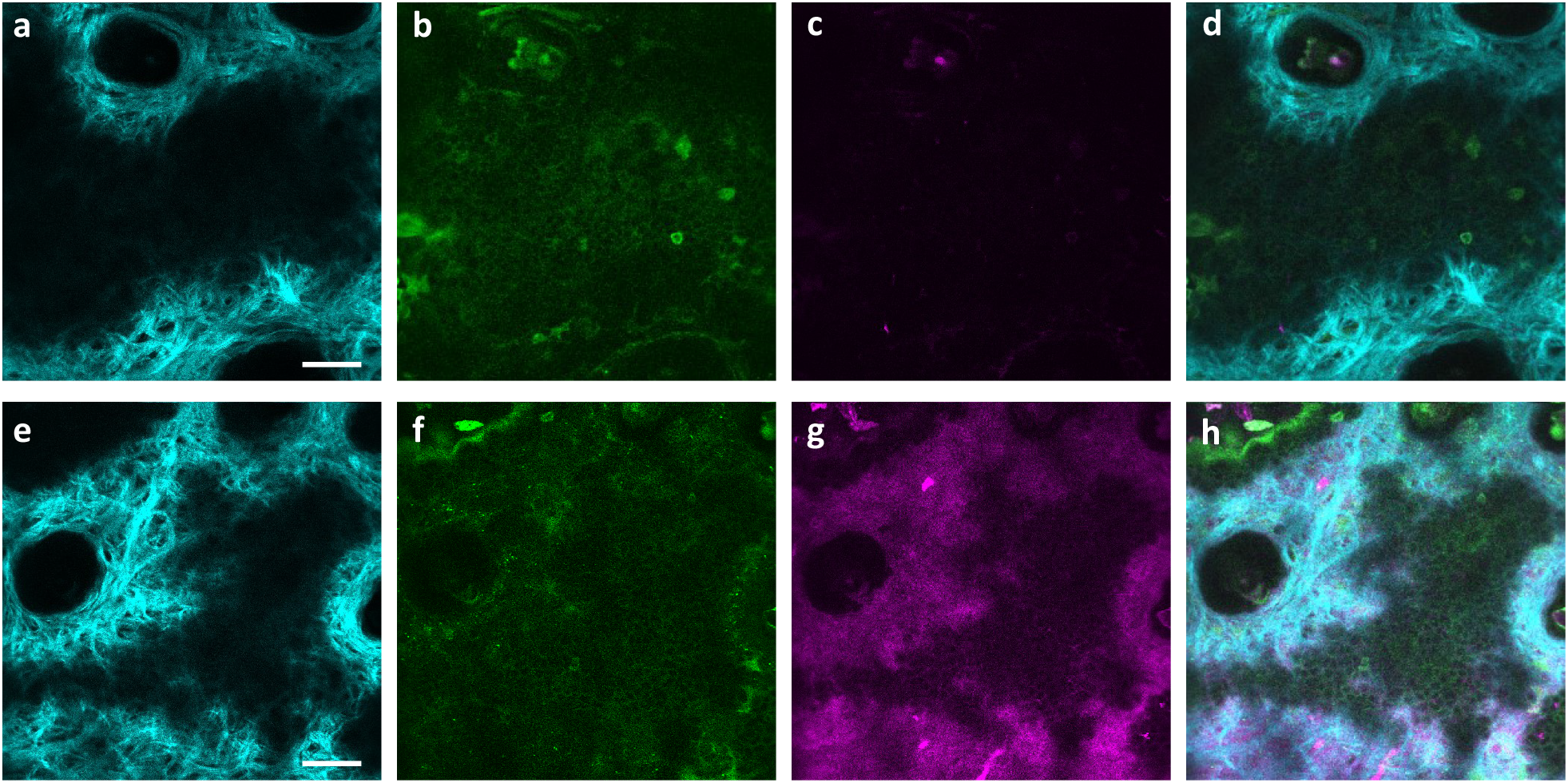
Intravital multiphoton microscopy of mouse skin. Images acquired with different emission channels of the multiphoton microscope while exciting the live mouse epidermis at 920 nm wavelength before (**a-d**) and after (**b-h**) injection of a high molecular weight fluorescent dextran. Second-harmonic generation (SHG) allows the imaging of collagen fibers, represented in cyan color (**a, e**); GCaMP6s and keratinocyte autofluorescence are shown in green color (**b, f**); fluorescent dextran is represented in magenta color; (**d, h**) composite images of the three channels. Basal keratinocytes (smaller polygonal cells in green) can be seen in the large flat area where collagen fibers are not present; larger polygonal cells are keratinocytes of the higher dermal layers. Oval-shaped spaces where both collagen fibers and keratinocytes are missing correspond to hair bulbs and related appendages, such as sebaceous glands. Scalebar: 50 µm.

### Image processing and data analysis

To establish a relationship between the [Ca^2+^]_C_ and the output signal of GCaMP6s, we treated GCaMP6s as a Ca^2+^ buffer (denoted as G) that is present in the cytosol at a concentration [G]. Chelation of Ca^2+^ by buffer G to form a complex CaG is described by the reaction:

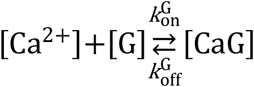

and the corresponding kinetic equation:

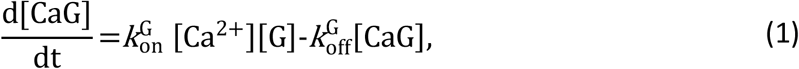

where square brackets are used to indicate concentration, 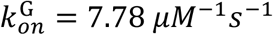 is the rate constant for Ca^2+^ binding to G and 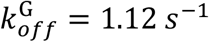 is the rate constant for Ca^2+^ dissociation [36].

At chemical equilibrium *d* [CaG]/*dt* = 0, therefore

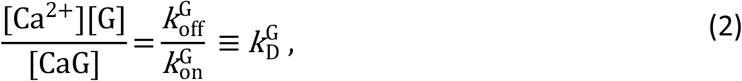

where 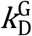 is the equilibrium or dissociation constant, thus

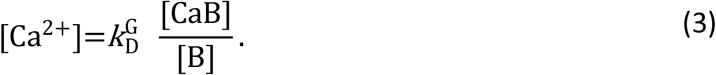

Assuming that [G] is low enough that the relationship between the fluorescence emission *F* (of GCaMP6s) and [G] is linear, *F* can be written as a linear combination:

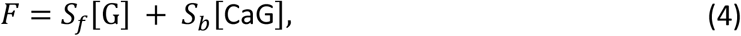

where the proportionality constant *S*_*f*_ and *S*_*b*_ lump all factors such as the illumination intensity, the efficiency of fluorescence emission, the molar absorption coefficient, and the length of the absorbing medium traversed by the illuminating beam. If *c*_*T*_ denotes the total (constant) fluorophore concentration, the mass balance equation is:

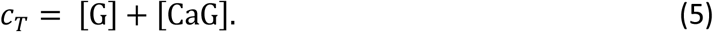

Defining the emission under Ca^2+^-saturating and Ca^2+^-free conditions respectively as *F*_*max*_ = *S*_*b*_*c*_*T*_ and *F*_*min*_ = *S*_*f*_*c*_*T*_, the expression for *F* can be re-written as

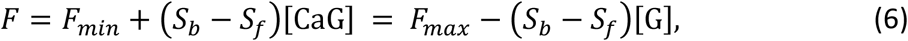

yielding

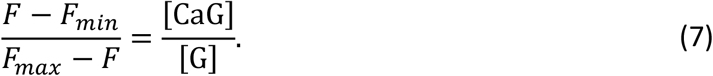

At chemical equilibrium, equation (2) holds. Therefore, we conclude that

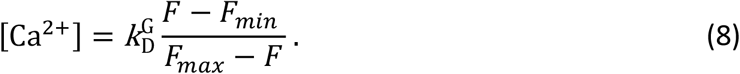

Equation (8) can be used to estimate the change in the cytosolic free Ca^2+^ concentration, Δ [Ca^2+^]_*c*_ *≡* [Ca^2+^]_*c*_ − [Ca^2+^]_0_, where [Ca^2+^]_0_ is the pre-stimulus (baseline) concentration. For changes that fall within the approximately linear region of the input/output relation, we may write:

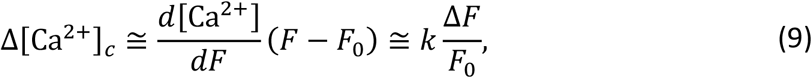

where

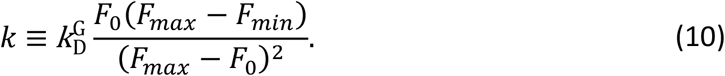

Thus, if imaging experiments are performed in such a way that bleaching is negligible, and as long *c*_*T*_ and the optical path length do not change during the measurement, the pixel-by-pixel ratio

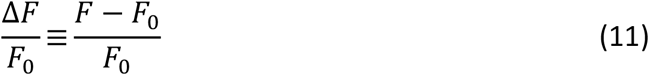

is a unique function of *Δ* [Ca^2+^]_C_ [39].

On this ground, image processing and data analysis used to relate *Δ* [Ca^2+^]_C_ and *F* were carried out using the open source ImageJ software, the MATLAB programming environment (R2015a, The MathWorks, Inc., Natick, MA, USA) and Vimmaging, a custom-made software developed under the MATLAB environment by Catalin D. Ciubotaru and Fabio Mammano.

All analyses were preceded by background subtraction in order eliminate contributions independent of GCaMP6s expression. To define the background-subtraction procedure, four images of basal layer keratinocytes were recorded from a wild type mouse (not expressing GCaMP6s, **Figure 4**) under the same conditions used for all other experiments (see **Table 1**). Histograms of pixel intensity were extracted from regions of interest (ROIs) of 76 × 76 pixels with ImageJ and fitted with a Gaussian distribution,

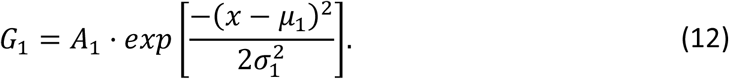

**Figure 4:**
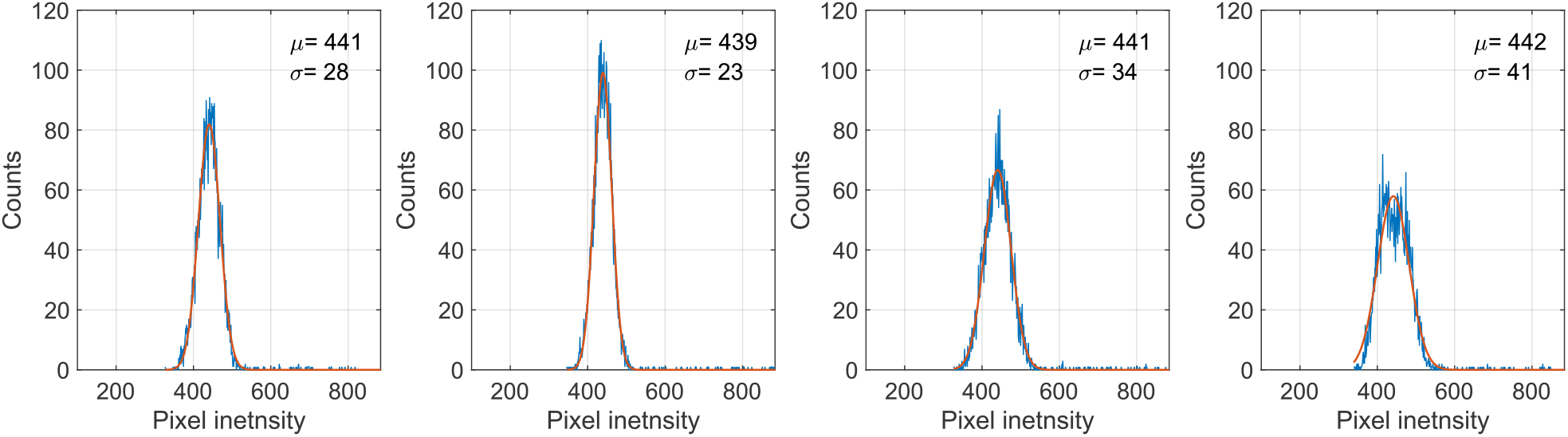
Distributions of background pixel intensity. Histograms of the pixel intensity (blue traces) in ROIs composed of 76 × 76 pixels within an image region where the keratinocytes were present. G_1_ fitting curves (orange traces) are shown together with the corresponding fitting paramaters, μ_1_ and σ _1_.

The fitting parameters for wild type mouse epidermis were *μ*_1_ = 440 ± 1 and *σ*_1_ = 31 ± 4.

Next, we examined basal layer keratinocytes in *n* = 15 GCaMP6s-expressing mice. The distribution of baseline pixel intensity values in ROIs composed of 76 × 76 pixels, averaged over the first 5 frames preceding the photodamage stimulus (that occurred at frame 6), followed a bimodal distribution that was well fitted by the sum of two Gaussians, *G*_1_ + *G*_2_,where

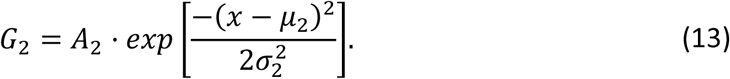

The parameters *μ*_1_ = 440 ± 2 and *σ*_1_ = 30 ± 1 (mean ± s.e.m) of *G*_1_ for the GCaMP6s samples were not significantly different from those of the wild type control (p = 0.98 for *μ*_1_; p = 0.5 for *σ*_1_ t-test). In contrast, the parameters *μ*_2_ = 697 ± 9 and *σ*_2_ = 206 ± 5 (mean ± s.e.m.) of *G*_2_ differed significantly from those of the wild type control (p = 2 × 10^10^ for *μ*_1_ ;p =2 × 10^12^ for *σ*_1_ t-test).Therefore, we attributed the pixel intensity distributions corresponding to *G*_1_ to the sum of instrument noise/offset plus keratinocyte autofluorescence, and the distributions corresponding to *G*_2_ to GCaMP6s pre-stimulus (baseline) fluorescence. Based on this analysis, the intersection between the two fitting curves (**Figure 5**, vertical dash-dotted lines), was selected as the threshold value to be subtracted from each image of the corresponding sequence. All pixel values below threshold were set to 1, thus eliminating autofluorescence and the instrumental offset. Thereafter, Ca^2+^ signals evoked by photodamage were quantified as the time-dependent background-subtracted pixel-by-pixel relative changes in fluorescence, Δ*F*(*t*)/*F*_0_, where *F*_0_ is the pre-stimulus fluorescence obtained by averaging over the first 5 (pre-stimulus) frames.

**Figure 5:**
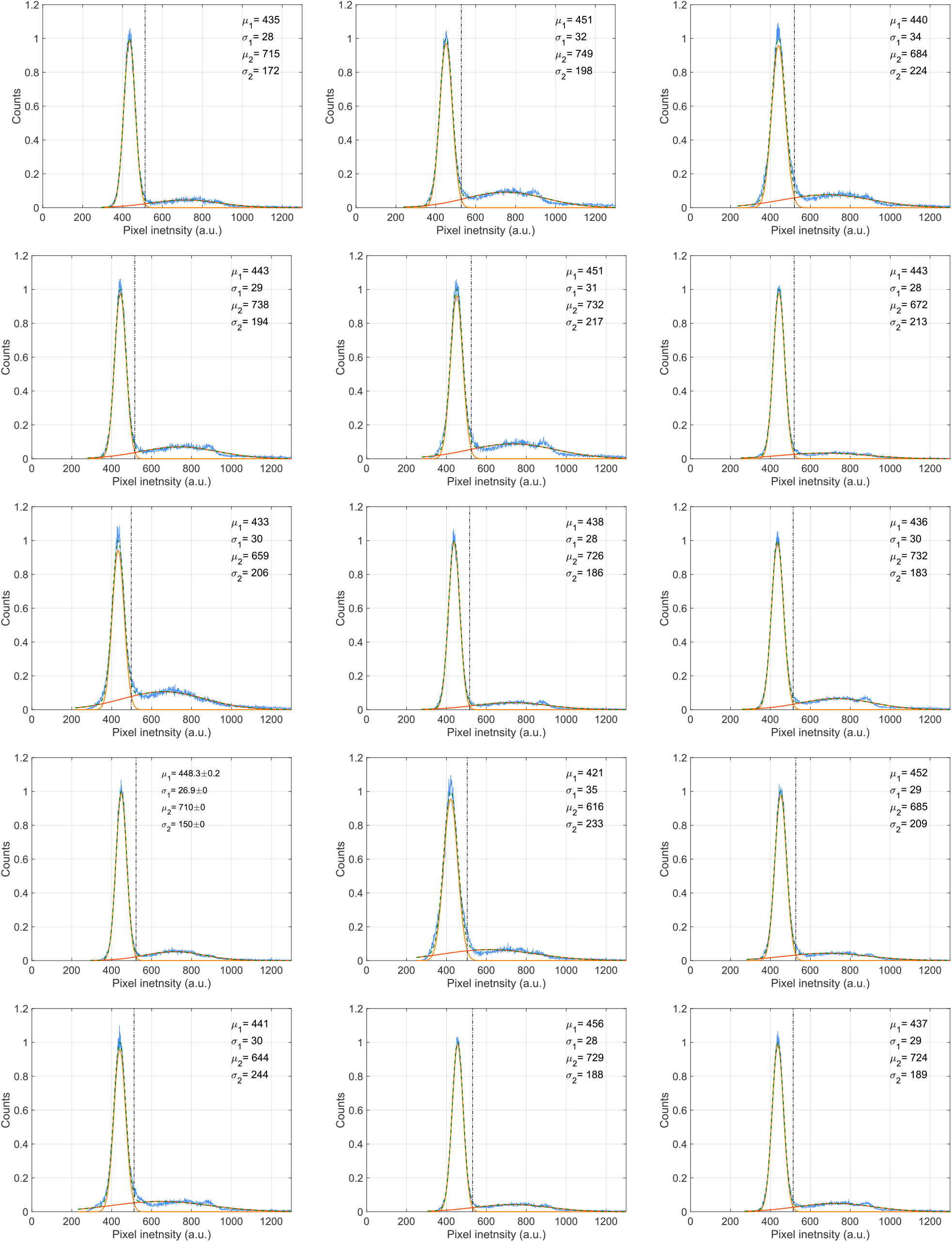
Background Subtraction. Shown are histograms of the pixel intensity (normalized to the maximum of the first peak) in a ROI of 76×76 pixels taken in a region where the keratinocytes were present in the first 5 frames before the photodamage. Each histogram corresponds to n ≥ 3 videos in m = 1 GCaMP6s expressing mouse (blue) and the double Gaussian fit (green dashed line) is shown together with the *μ* and *σ* of each gaussian distribution. The intersection between the two fitting distribtiuons (yellow and orange), marked by a vertical dash-dotted line, was used for background subtraction (see text).

### Statistical analysis

Comparisons of means for non-gaussian sample distributions were made using the Wilcoxon Rank sum test (as implemented in the Matlab function *ranksum*). For samples that had Gaussian distribution, the two-sample t-test (as implemented in the Matlab function *ttest2*) was adopted. P = p-value < 0.05 was assumed as statistically significant. Mean values are quoted ± standard error of the mean (s.e.m.).

## RESULTS

### Dynamics of Ca^2+^ waves elicited by focal intradermal photodamage in the earlobe skin of live anesthetized mice

To photodamage a single keratinocyte in the basal epidermal layer of earlobe skin in live anesthetized mice expressing the GCaMP6s Ca^2+^ indicator, we used the second scanning head of the multiphoton microscope (GG branch of the optical path, see Materials and methods). Ca^2+^ waves propagated radially from the damaged to the surrounding/bystander epidermal cells (referred to as expansion phase, **Figure 6 a**). Fluorescence signals persisted for several minutes in keratinocytes invaded by the Ca^2+^ wave, and slowly returned to basal levels (referred to as waning phase, **Figure 6 b**). Mean ± s.e.m. of the area *A*(*t*) invaded by the Ca^2+^ wave are shown for both expansion and waning phases in **Figure 6 c, d**. *A*(*t*) was computed by counting all pixels in which Δ*F*(*t*)/*F*_0_ (see Materials and Methods) exceeded an arbitrary threshold, corresponding to 30% of the maximum signal achieved as a result of photodamage in the given image sequence.

**Figure 6:**
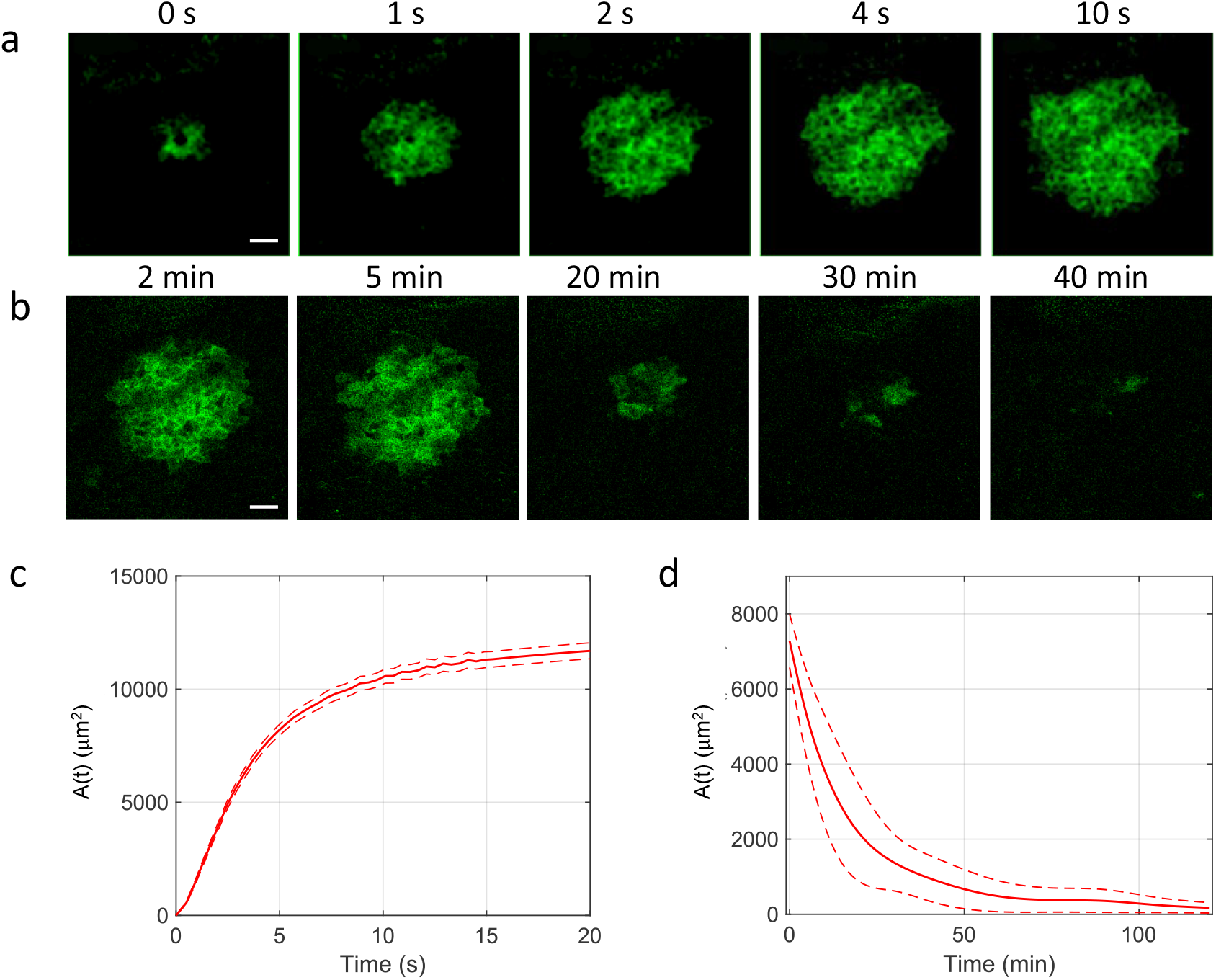
Intravital multiphoton microscopy of Ca^2+^ waves elicited by focal photodamage in the epidermal basal layer of earlobe skin in live anesthetized mice. (**a, b**) Representative sequences of GCaMP6s fluorescence *Δ F* images acquired at shown time points from the end of a 0.5 s photodamage interval (indicated as 0 s), showing Ca^2+^ wave expansion (**a**) and subsequent waning (**b**). Scale bars: 25 μm. (**c, d**) Invaded area vs. time, *A*(*t*), during Ca^2+^ wave expansion (**c**) and waning (**d**). (**c**) Mean (solid line) ± s.e.m. (dashed lines) of *n* = 60 experiments in *m* = 14 mice. (**d**) mean (solid line) ± s.e.m. (dashed lines) of *n* = 3 experiments in *m* = 1 mouse.

The volume of tissue invaded by the Ca^2+^ wave at the time of maximal expansion (*V*= 1.2 ± 0.2 ×10^5^ μm^3^) was visualized by acquiring through-focus image sequences (also known as *z*-stacks) at 2 µm increments along the optical axis (*z*) of the objective lens (**Figure 7 a, b**). The graph *A*(*z*) of the invaded area vs. depth (**Figure 7 c**) shows that Ca^2+^ waves extended up to ∼10 µm above the photodamage plane, and occasionally stimulated cells with the typical morphology of Langerhan cells, reaching the surface of the thin earlobe mouse skin (corneocyte layer). In the opposite, dermal direction, Ca^2+^ waves reached down to ∼20 µm below the photodamage plane invading the fibroblast-populated collagen matrix.

**Figure 7:**
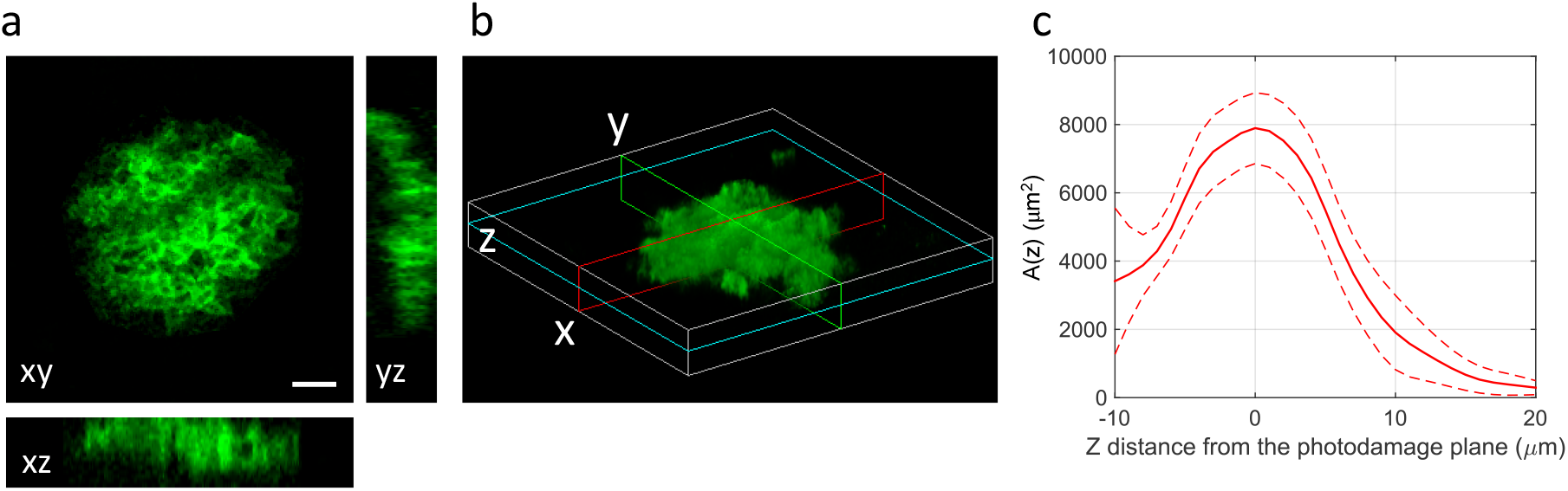
Volume of tissue invaded by the Ca^2+^ wave after focal photodamage. (**a**) Views of the wave in the xy plane (focal plane) and two orthogonal (xz and yz) planes. (**b**) 3D rendering of the volume invaded by the wave; cyan, red, green and lines indicate the xy, xz and yz planes of panel (**a**), respectively. (**c**) Invaded area vs. axial coordinate (z, along the optical axis of the objective lens); data are mean (solid line) ± s.e.m. (dashed lines) of n = 7 experiments in m = 1 mouse. Scale bar: 25 µm.

Given the approximate radial symmetry of Ca^2+^ waves during the expansion phase in the focal plane, the equivalent radius of the invaded area was computed as

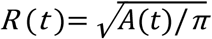

(**Figure 8 a**). The maximal value *R*_max_= max [*R* (*t*)] averaged over the control experiments that gave the largest waves was 67 ± 1 µm (mean ± s.e.m. ; *n* = 21 experiments in *m* = 6 mice).

**Figure 8:**
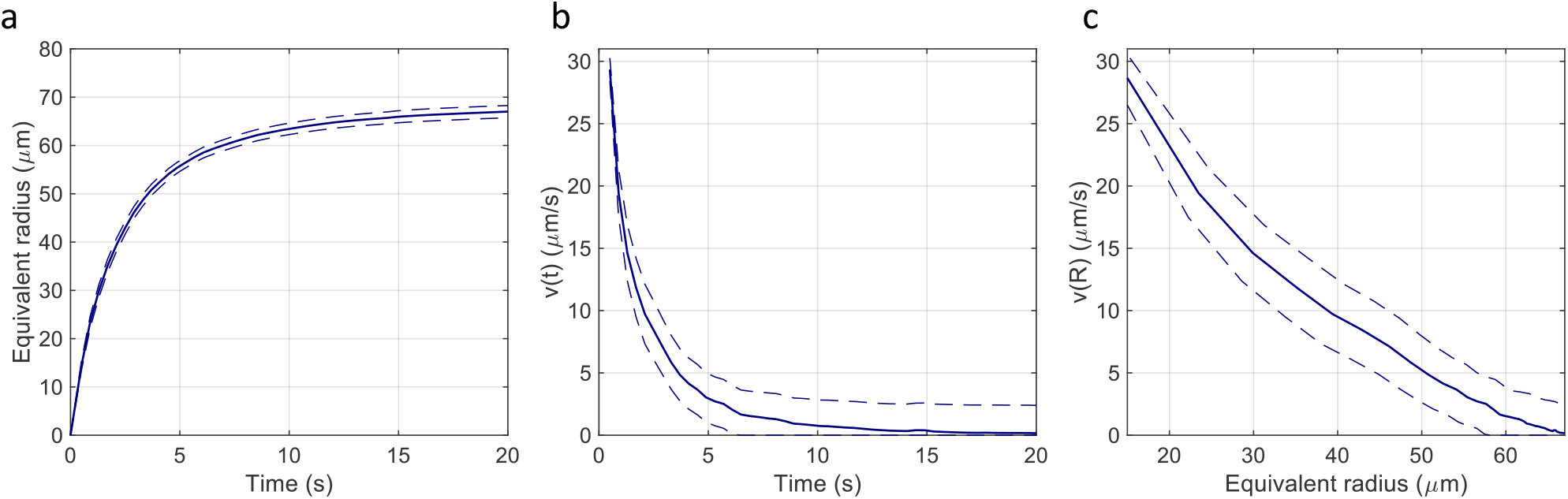
Quantitative analysis of the expansion phase. (**a**) Equivalent radius of the invaded area during Ca^2+^ wave expansion; propagation velocity of the Ca^2+^ wave as function of time (**b**) and of the equivalent radius (**c**). Mean (blue solid lines) ± s.e.m. (blue dashed lines) of *n*=21 experiments in *m*=6 mice.

The propagation velocity of the wave during the expansion phase was computed as time derivative of the equivalent radius

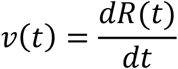

and it was evaluated as function of time and of the equivalent radius *R* (*t*) (**Figure 8 b**,**c**), both displaying a monotonical decrease towards zero.

To further characterize the expansion phase, we computed whole-cell Δ*F*(*t*)/*F*_0_ responses by spatially averaging pixel signals over individual bystander keratinocytes. Data from different keratinocytes in different experiments were grouped and averaged based on the distance of the cell centroid from the photodamage site in the range from 10 ± 3 µm to 80 ± 3 µm (**Figure 9 a**). Note that this range exceeds the average value of *R*_max_ given above. The discrepancy is due to the 30% threshold criterion used to select pixels that contributed to the estimate of the area *A*(*t*) invaded by the Ca^2+^ wave. In general Δ*F*(*t*)/*F*_0_ responses at 70 ± 3 µm and 80 ± 3 µm remained subthreshold, and therefore did not contribute to *A*(*t*).

**Figure 9:**
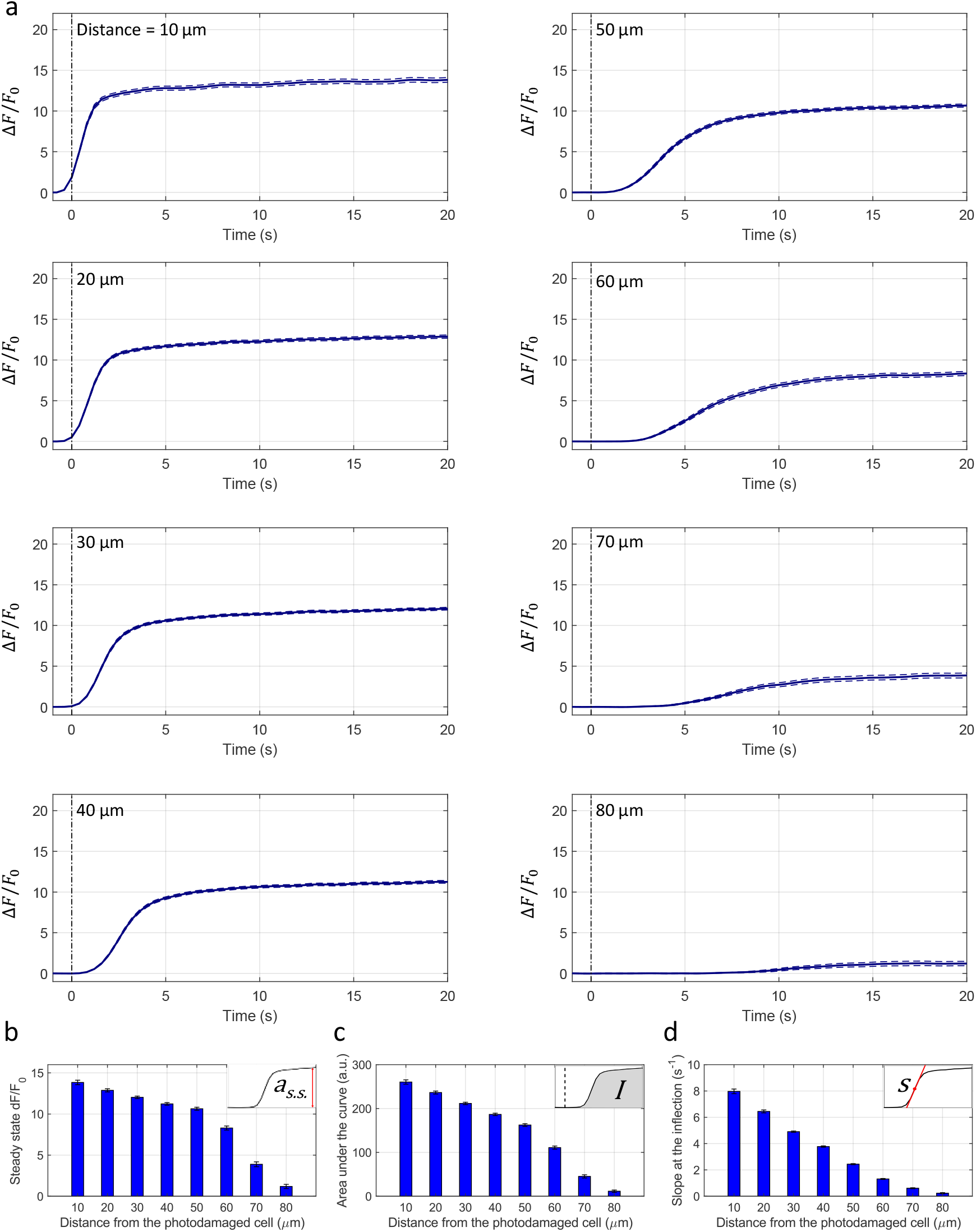
Analysis of GCaMP6s fluorescence changes in bystander keratinocytes at increasing distance from the photodamage site. (**a**) In each panel, the vertical black dash-dotted line at 0 s marks the end of the 0.5 s photodamage time interval. Data are mean (solid line) ± s.e.m. (dashed lines) vs. time for n = 60 experiments in m = 15 mice. (**b**) Amplitude (*a*_*s*.*s*._) of the Δ*F*(*t*)/*F*_0_ signal at steady state. (**c**) Area (*I*) under the Δ*F*(*t*)/*F*_0_ trace, computed between 0 and 20 s. (**d**) Slope (*S*) of the Δ*F*(*t*)/*F*_0_ trace at the inflection point (mean ± s.e.m). Data in (**b-d**) are mean ± s.e.m. vs. bystander cell distance from the photodamage site.

From the analysis of Δ*F*(*t*)/*F*_0_ traces, we extracted 3 parameters for each order of bystander keratinocytes. The first parameter is the steady state amplitude

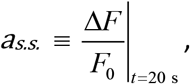

which decreased slowly with distance *d* from the photodamage site, up to a critical value *d*_stop_(typically comprised between 70 µm and 80 µm) beyond which wave propagation ceased abruptly and *a*_*s*.*s*_ approached zero (**Figure 9 b**).

The second parameter is the Ca^2+^ load imparted on the cell by the Ca^2+^ wave, computed as the integral

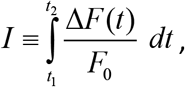

where *t*_1_ = 0 and *t*_2_ = 20 s. *I* increases if the amplitude of Δ*F*(*t*)/*F*_0_ increases and/or the velocity of wave propagation increases (due to the fixed limits of the integration interval). For the same reason, due to the progressively delayed rise of the Ca^2+^ response, *I* decreased with the distance *d* from the photodamage site (**Figure 9 c**).

The third parameter is the response speed of the cell

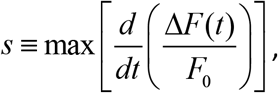

computed as the slope of the trace around the inflection point of its rising phase. Clearly, *S* depends both on the velocity of wave propagation and the kinetics of the signal transduction chain that promotes the [Ca^2+^]_C_ increase in the given cell. The results show that *S* was a monotonically decreasing function of the distance *d* from the photodamage site (**Figure 9 d**).

To explore the mechanisms underlying the propagation of Ca^2+^ waves, we used different drugs that interfere (more or less specifically) with candidate pathways. Waves evoked in control conditions were compared with those obtained after drug delivery in the same animal. As a rule, drug-conditioned waves were evoked in and propagated through naïve epidermal locations, i.e. areas of the epidermis not reached by control waves. Initially, a dedicated set of experiments were conducted to ascertain the effects of microinjecting solely vehicle solution (VS = PBS supplemented with Dextran Texas Red, 1.78 μM; see Materials and Methods) in the earlobe skin proximal to the photodamage site. Statistical analyses of the results indicate that the parameters used to characterize wave expansion after photodamage were largely unaffected by VS microinjection (**Figure 10**). This experiment was repeated in other 2 mice, with similar results.

**Figure 10:**
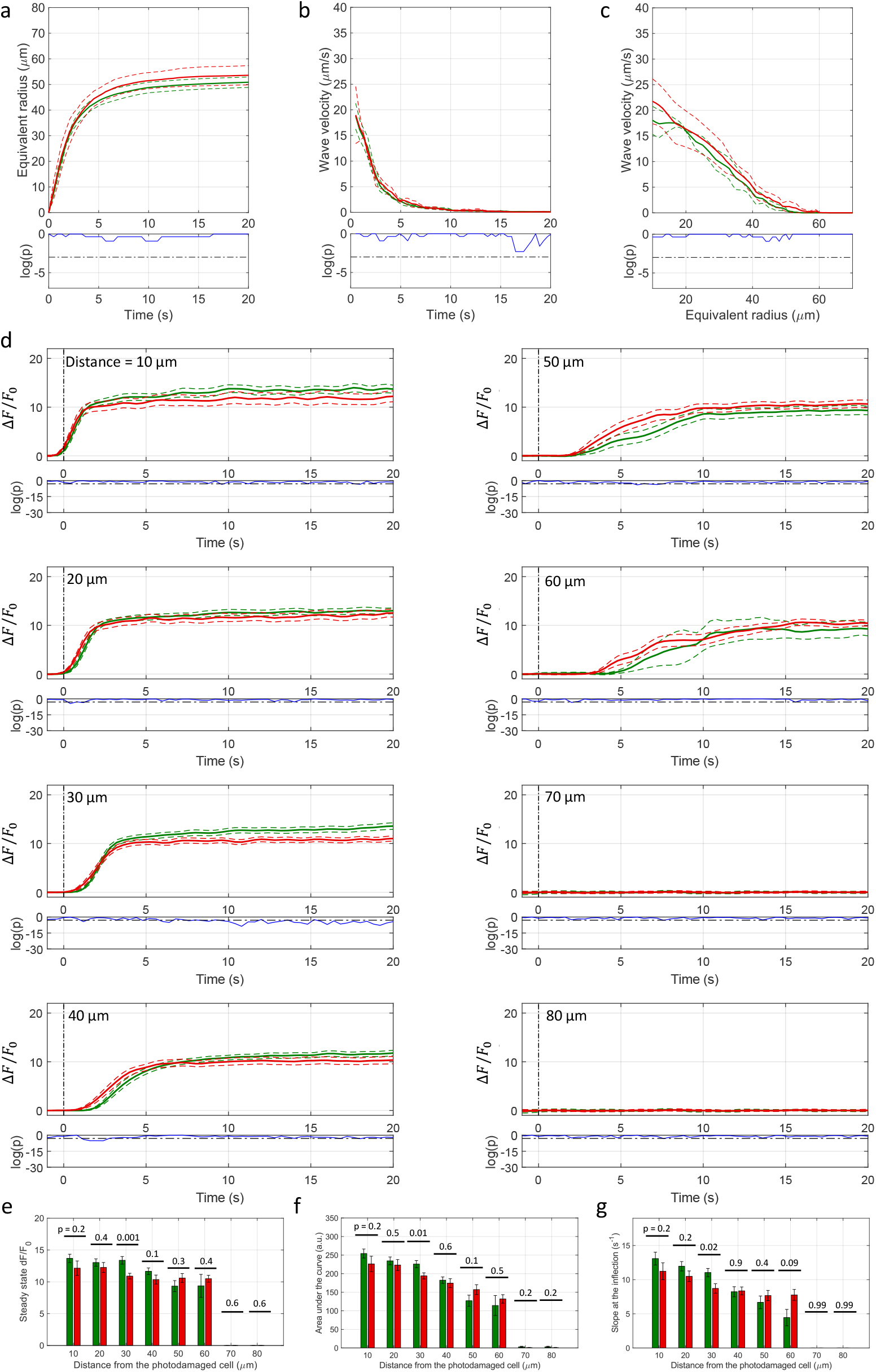
Effect of VS microinjection on Ca^2+^ wave expansion after focal photodamage. (**a**) Equivalent radius of the area invaded by Ca^2+^ waves as a function of time after photodamage; speed of the expanding wave as function of time (**b**) and of the equivalent radius (**c**). (**d**) Δ*F*(*t*)/*F*_0_ responses of bystander keratinocytes at increasing distance from the photodamage site. In each panel, the vertical black dash-dotted line at 0 s marks the end of the 0.5 s photodamage time interval. Data in (**a-d**) are mean (solid line) ± s.e.m. (dashed line) in control conditions (green) and after VS microinjection (red). Point-by-point p-values (p; Wilcoxon Rank Sum test for **a-c**; two-sample t-test for **d**) are shown on a logarithmic scale below each graph (blue traces); p < 0.05 (horizontal black dash-dotted line) indicates statistical significance. (**e**) Amplitude (*a*_*s*.*s*._) of the Δ*F*(*t*)/*F*_0_ trace at steady state. (**f**) Area (*I*) under the Δ*F*(*t*)/*F*_0_ trace, computed between 0 and 20 s. (**g**) Slope (*S*) of the Δ*F*(*t*)/*F*_0_ trace at the inflection point. Data in (**e-g**) are mean ± s.e.m. vs. bystander cell distance from the photodamage site in control conditions (green) and after VS microinjection (red). P-values are shown above each pair of bars (two-sample t-test).

Because VS injection had no significant influence on the parameters used to characterize quantitatively the expansion phase of Ca^2+^ waves, all successive experiments were performed in the following way: (1) a certain number of waves (used as reference) were evoked in non-overlapping areas of the naïve mouse earlobe epidermis; thereafter, (2) the drug of interest dissolved in VS was microinjected intradermally (3) and second set of waves were evoked (in non-overlapping areas reached by Dextran Texas Red), (4) recorded and (5) compared with the previously acquired reference waves.

### Role of extracellular ATP

ATP in the extracellular milieu may couple to P2 purinoceptors on the cell plasma membrane to develop intercellular Ca^2+^ waves with or without involvement of IP_3_ diffusion through intercellular gap junction channels (IGJCs) [31]. To test the hypothesis that photodamage triggered intercellular Ca^2+^ waves by promoting ATP release from the photodamaged keratinocyte, we injected 4 µl of VS containing apyrase (500 U/mL), an enzyme that catalyzes ATP hydrolysis [40]. The presence of microinjected apyrase in the extracellular milieu correlated with a highly significant reduction of the invaded area *A*(*t*), therefore also of the equivalent Ca^2+^ wave radius *R* (*t*) (**Figure 11 a**). From the same experiments, we derived and analyzed Δ*F*(*t*)/*F*_0_ traces from bystander keratinocytes and confirmed that apyrase severely hampered Ca^2+^ wave propagation, such that keratinocytes at distances *d* > 20 μm from the photodamage site failed to respond (**Figure 11 d**). Consequently, all three critical parameters used to characterize these responses (a_s.s._, *I* and s) were null at *d* > 20 μm (**Figure 11 e-g**). These experiments were conducted in *n*=4 (control) and *n*=4 (apyrase) non-overlapping areas of earlobe skin in *m*=1 mouse.

**Figure 11:**
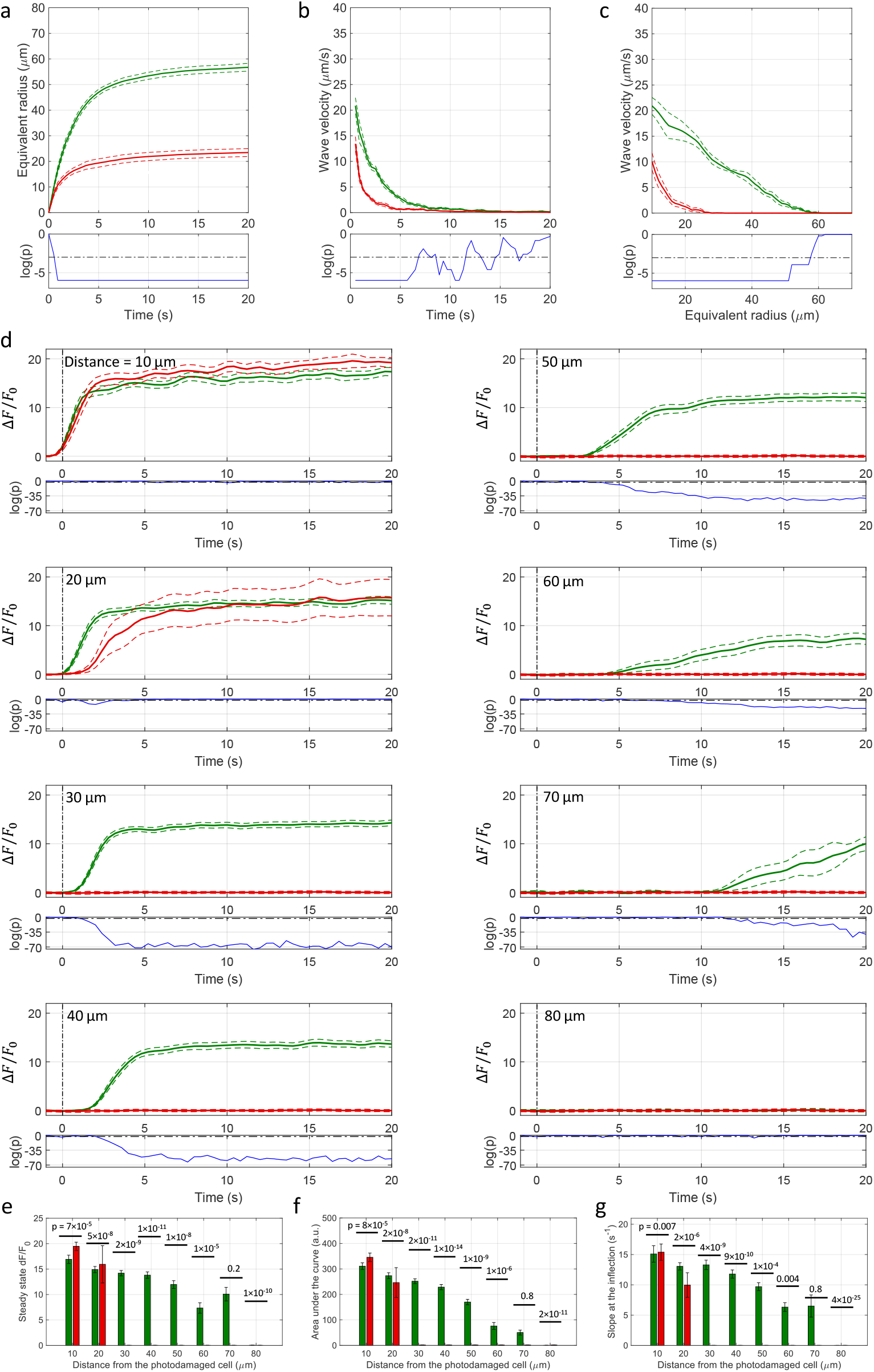
Effect of apyrase. (**a**) Equivalent radius of the area invaded by Ca^2+^ waves as a function of time after photodamage; speed of the expanding wave as function of time (**b**) and of the equivalent radius (**c**). (**d**) Δ*F*(*t* /*F*_0_ responses of bystander keratinocytes at increasing distance from the photodamage site. In each panel, the vertical black dash-dotted line at 0 s marks the end of the 0.5 s photodamage time interval. Data in (**a-d**) are mean (solid line) ± s.e.m. (dashed line) in control conditions (green) and after 500 U/ml apyrase microinjection (red). Point-by-point p-values (p; Wilcoxon Rank Sum test for **a-c**; two-sample t-test for **d**) are shown on a logarithmic scale below each graph (blue traces); p < 0.05 (horizontal black dash-dotted line) indicates statistical significance. (**e**) Amplitude (*a*_*s*.*s*._) of the Δ*F*(*t*)/*F*_0_ signal at steady state. (**f**) Area (*I*) under the Δ*F*(*t*)/*F*_0_ trace, computed between 0 and 20 s. (**g**) Slope (*S*) of the Δ*F*(*t*)/*F*_0_ trace at the inflection point. Data in (**e-g**) are mean ± s.e.m. vs. bystander cell distance from the photodamage site in control conditions (green) and after apyrase microinjection (red). P-values are shown above each pair of bars (two-sample t-test).

To determine whether P2 purinoceptors contributed to photodamage-evoked Ca^2+^ waves, we microinjected 4 µl of VS containing pyridoxal phosphate-6-azo(benzene-2,4-disulfonic acid) tetrasodium salt hydrate (PPADS, 625 µM), a P2 purinoceptor antagonist [24, 41]. PPADS reduced significantly the area invaded by Ca^2+^ waves following focal photodamage (**Figure 12 a**) and slowed down significantly Δ*F*(*t*)/*F*_0_ responses in all bystander keratinocytes (**Figure 12 b-d**). The parameters a_*s*.*s*._, *I* and S were significantly reduced in PPADS compared to controls at *d* > 30 μm, and the arrest distance *d*_stop_ decreased from 80 μm (control) to 50 μm (PPADS) (**Figure 12 e-g**). These experiments were conducted in *n*=4 (control) and *n*=5 (PPADS) non-overlapping areas of earlobe skin in *m*=1 mouse.

**Figure 12:**
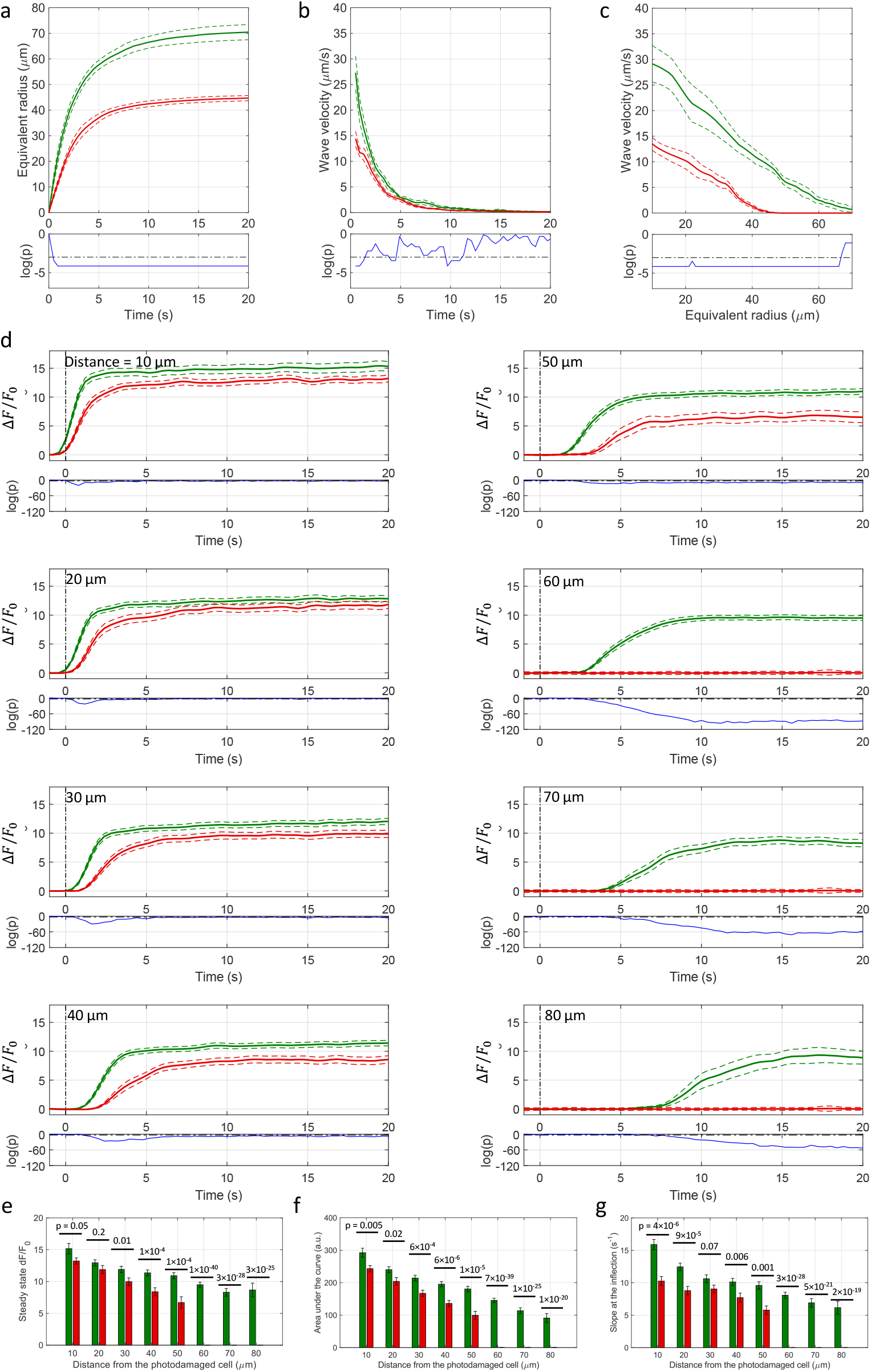
Effect of PPADS. (**a**) Equivalent radius of the area invaded by Ca^2+^ waves as a function of time after photodamage; speed of the expanding wave as function of time (**b**) and of the equivalent radius (**c**). (**d**) Δ*F*(*t*)/*F*_0_ responses of bystander keratinocytes at increasing distance from the photodamage site. In each panel, the vertical black dash-dotted line at 0 s marks the end of the 0.5 s photodamage time interval. Data in (**a-d**) are mean (solid line) ± s.e.m. (dashed line) in control conditions (green) and after PPADS (625 µM) microinjection (red). Point-by-point p-values (p; Wilcoxon Rank Sum test for **a-c**; two-sample t-test for **d**) are shown on a logarithmic scale below each graph (blue traces); p < 0.05 (horizontal black dash-dotted line) indicates statistical significance. (**e**) Amplitude (*a*_*s*.*s*._) of the Δ*F*(*t*)/*F*_0_ signal at steady state. (**f**) Area (*I*) under the Δ*F*(*t*)/*F*_0_ trace, computed between 0 and 20 s. (**g**) Slope (*S*) of the Δ*F*(*t*)/*F*_0_ trace at the inflection point. Data in (**e-g**) are mean ± s.e.m. vs. bystander cell distance from the photodamage site in control conditions (green) and after PPADS microinjection (red). P-values are shown above each pair of bars (two-sample t-test).

Once released in the extracellular space, ATP is rapidly hydrolyzed by cell surface-located ectonucleotidases, in particular by members of the ecto-nucleoside triphosphate diphosphohydrolase family (NTPDase1,2,3, and 8) [42-44]. Therefore, we predicted that interfering with ectonucleotidase function should influence Ca^2+^ wave propagation. To test this hypothesis, we injected 4 µl of VS containing ARL 67156 trisodium salt hydrate (shortened as ALR, 400 µM), a widely used NTPDase inhibitor [45-47]. The presence of microinject ARL in the extracellular milieu correlated with a significant increase of the area *A*(*t*) invaded by Ca^2+^ waves, hence of *R* (*t*) (**Figure 13 a**). ARL increased also the velocity *V* (*t*) of Ca^2+^ wave propagation, confirmed by the significantly reduced lag with which Δ*F*(*t*)/*F*_0_ traces grew after photodamage in bystander keratinocytes at distances in excess of 20 μm (**Figure 13 b-d**). In addition, ARL increased significantly the parameters *a*_*s*.*s*._, *I* and *S* at distances in excess of 40 μm (**Figure 13 e-g**). These experiments were conducted in *n*=4 (control) and *n*=4 (ARL) non-overlapping areas of earlobe skin in *m*=1 mouse.

**Figure 13:**
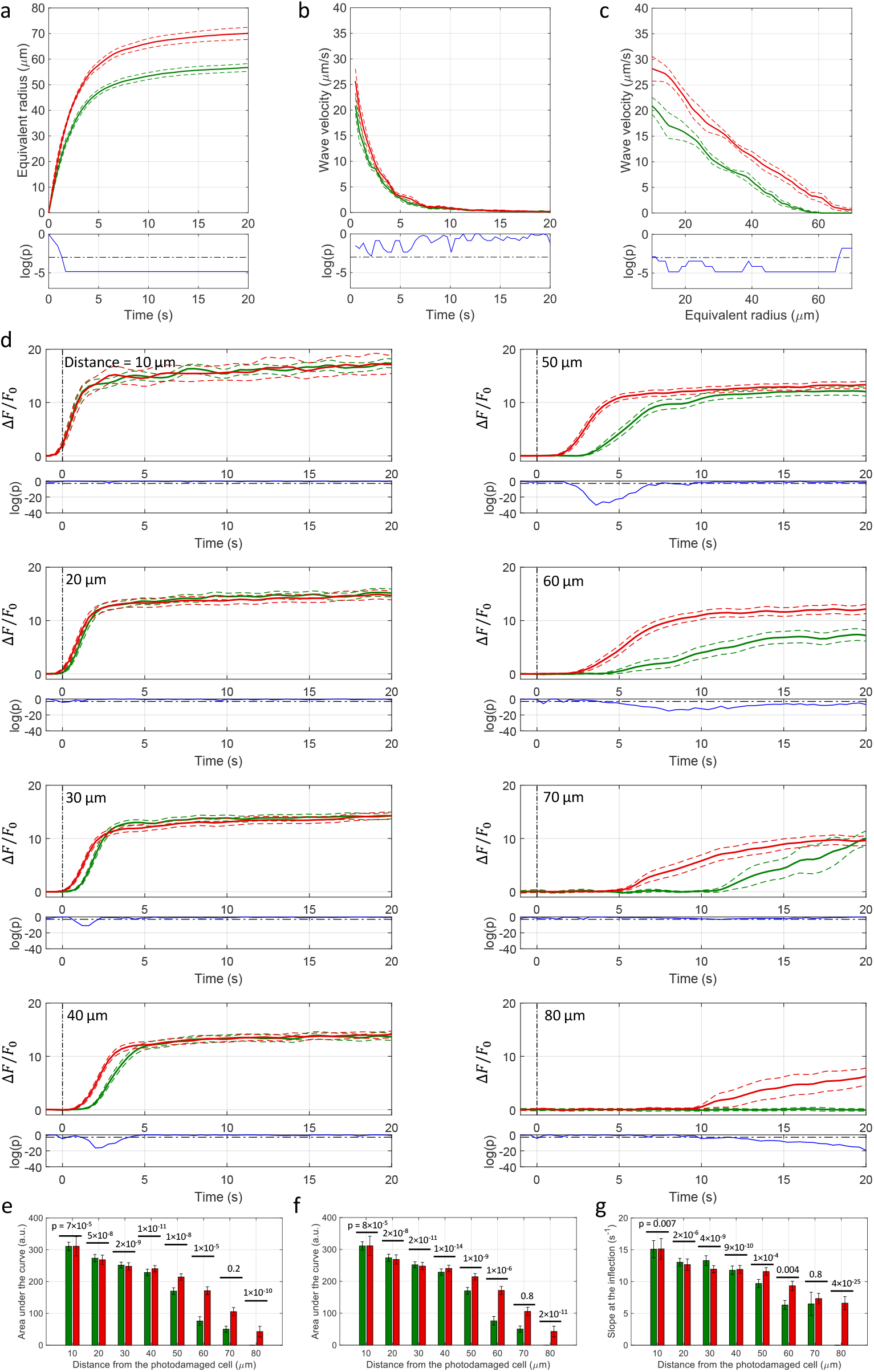
Effect of ARL. (**a**) Equivalent radius of the area invaded by Ca^2+^ waves as a function of time after photodamage; speed of the expanding wave as function of time (**b**) and of the equivalent radius (**c**). (**d**) Δ*F*(*t*)/*F*_0_ responses of bystander keratinocytes at increasing distance from the photodamage site. In each panel, the vertical black dash-dotted line at 0 s marks the end of the 0.5 s photodamage time interval. Data in (**a-d**) are mean (solid line) ± s.e.m. (dashed line) in control conditions (green) and after ARL (400 µM) microinjection (red). Point-by-point p-values (p; Wilcoxon Rank Sum test for **a-c**; two-sample t-test for **d**) are shown on a logarithmic scale below each graph (blue traces); p < 0.05 (horizontal black dash-dotted line) indicates statistical significance. (**e**) Amplitude (*a*_*s*.*s*._) of the Δ*F*(*t*)/*F*_0_ signal at steady state. (**f**) Area (*I*) under the Δ*F*(*t*) *F* /*F*_0_ trace, computed between 0 and 20 s. (**g**) Slope (*S*) of the Δ*F*(*t*)/*F*_0_ trace at the inflection point. Data in (**e-g**) are mean ± s.e.m. vs. bystander cell distance from the photodamage site in control conditions (green) and after ARL microinjection (red). P-values are shown above each pair of bars (two-sample t-test).

Together, the results in **Figures 11-13** indicate that (ii) extracellular ATP is a key signaling molecule underlying Ca^2+^ wave propagation in the photodamaged epidermis; (ii) the area *A*(*t*) invaded by the Ca^2+^ wave and its velocity of propagation *V*(*t*) depend on degradation of extracellular ATP by NTPDases expressed by epidermal keratinocytes (which can be partially inhibited by ARL).

### Role of extracellular Ca^2+^

To determine the role of extracellular Ca^2+^ in the dynamics of photodamage-evoked Ca^2+^ waves, we microinjected 4 µl of VS containing 5 mM EGTA, a widely used Ca^2+^ chelator [48]. Reduction of the extracellular free Ca^2+^ concentration ([Ca^2+^]_ex_) with EGTA caused a paradoxical enhancement of the invaded area, gauged by the effective radius *R* (*t*) (**Figure 14 a**), paralleled by a significant increase of Δ*F*(*t*)/*F*_0_ bystander responses at all distances within 60 μm from the photodamaged cell (**Figure 14 d**). The parameters a_*s*.*s*._, *I* and S were significantly increased in EGTA compared to controls, and the arrest distance *d*_stop_ increased from 70 μm (control) to 80 μm (EGTA) (**Figure 14 e-g**). These experiments were conducted in *n*=5 (control) and *n*=6 (EGTA) non-overlapping areas of earlobe skin in *m*=1 mouse and have important implications for our understanding of Ca^2+^ dynamics.

**Figure 14:**
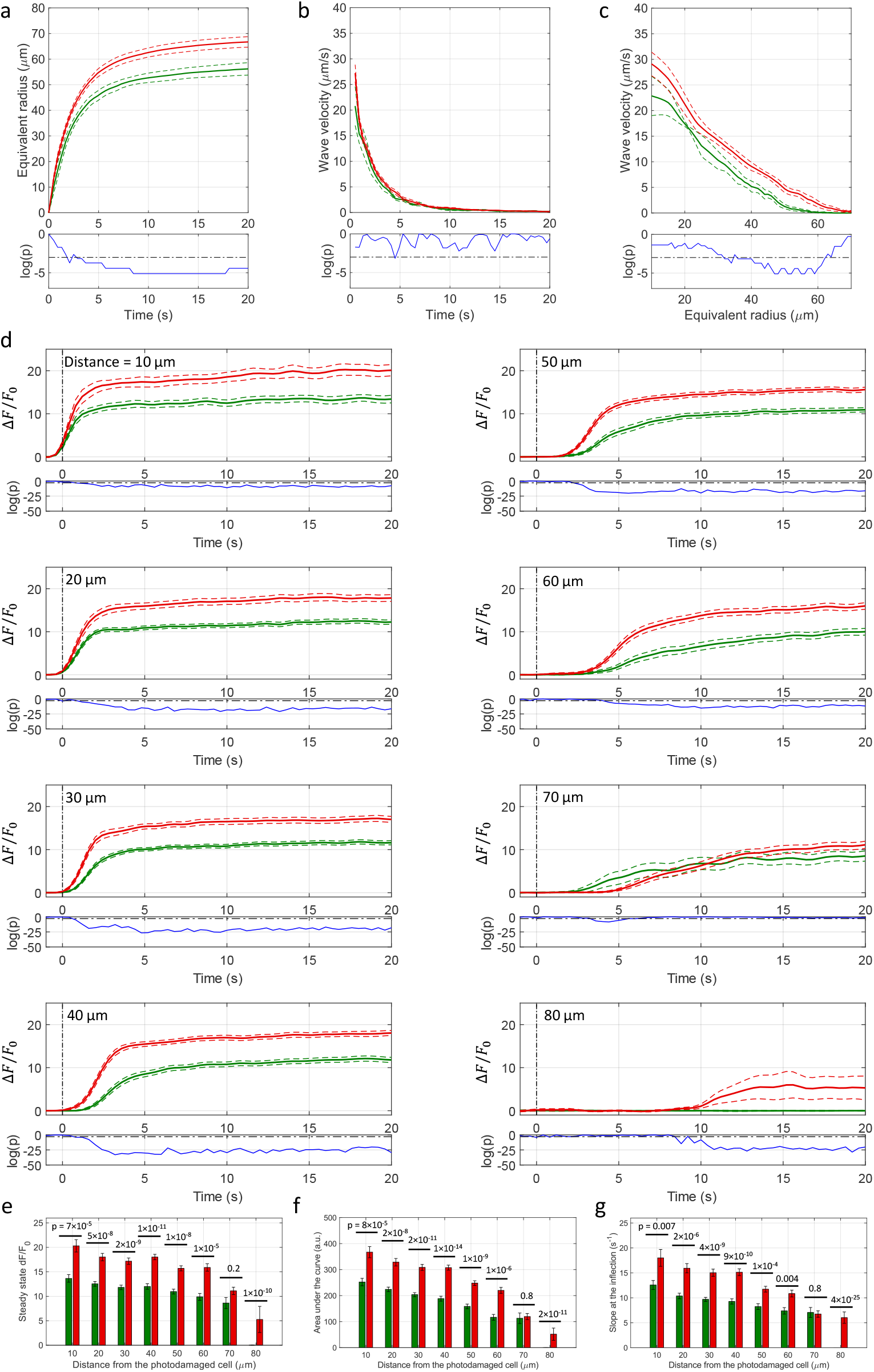
Effect of EGTA. (**a**) Equivalent radius of the area invaded by Ca^2+^ waves as a function of time after photodamage; speed of the expanding wave as function of time (**b**) and of the equivalent radius (**c**). (**d**) Δ*F*(*t*)/*F*_0_ responses of bystander keratinocytes at increasing distance from the photodamage site. In each panel, the vertical black dash-dotted line at 0 s marks the end of the 0.5 s photodamage time interval. Data in (**a-d**) are mean (solid line) ± s.e.m. (dashed line) in control conditions (green) and after EGTA (5 mM) microinjection (red). Point-by-point p-values (p; Wilcoxon Rank Sum test for **a-c**; two-sample t-test for **d**) are shown on a logarithmic scale below each graph (blue traces); p < 0.05 (horizontal black dash-dotted line) indicates statistical significance. (**e**) Amplitude (*a*_*s*.*s*._) of the Δ*F*(*t*)/*F*_0_ signal at steady state. (**f**) Area (*I*) under the Δ*F*(*t*)/*F*_0_ trace, computed between 0 and 20 s. (**g**) Slope (*S*) of the Δ*F*(*t*)/*F*_0_ trace at the inflection point. Data in (**e-g**) are mean ± s.e.m. vs. bystander cell distance from the photodamage site in control conditions (green) and after EGTA microinjection (green). P-values are shown above each pair of bars (two-sample t-test).

First, as EGTA increased Ca^2+^ responses, Ca^2+^ influx through plasma membrane channels (including P2X receptors [21]) did not to contribute significantly to the observed increments of the [Ca^2+^]_c_. Hence, the latter should be attributed chiefly to signal transduction through P2Y receptors (P2YR), via the coupling to G proteins, activation of phospholipase C (PLC), leading to the formation of IP_3_ and release of Ca^2+^ from intracellular stores [22-24, 49]. PPADS antagonizes P2Y_1_R [50], P2Y_4_R [51], P2Y_6_R [52] and P2Y_13_ R[53] (see also ref. [22]). In contrast, P2Y_2_R is PPADS-insensitive [22, 51]. Analysis of P2Y mRNA expression in murine keratinocytes identified P2Y_1_R, P2Y_2_R, P2Y_4_R and P2Y_6_R subtypes [54], therefore P2Y_2_R may be responsible for the incomplete effect of PPADS administration in our photodamage experiments (**Figure 12**).

Second, human and murine NTPDases require Mg^2+^ or Ca^2+^ ions for their activity, with optimal concentration between 1 and 5 mM [44]. Therefore, similar to ARL, also EGTA might indirectly inhibit ATP degradation leading to Ca^2+^ wave enhancement (**Figure 13 a**). In addition, lowering the [Ca^2+^]_ex_ with EGTA favors the opening of connexin hemichannels (HCs) [55]. Open HCs mediate the diffusive release of paracrine messengers, most importantly ATP [56-60]. Various expression profiles of connexins through epidermis layers are reported in literature, the majority of them agreeing in the predominance of Cx43 in the basal layer of the epidermis [61-72]. Thus, if present, HCs in basal keratinocytes are most likely to be composed of Cx43 protomers. Also pannexin 1 (Panx1) channels [73] are permeable to ATP [74], expressed in the epidermis and important for differentiation of keratinocytes and wound healing [75]. In addition, Panx1 channels can be activated by extracellular ATP acting through purinergic receptors of the P2Y group as well as by cytoplasmic Ca^2+^ [76]. Therefore, a positive-feedback loop involving the control by cytoplasmic Ca^2+^ of Panx1 [76] and/or connexin HCs [77-79] may lead to an overall augmented ATP-induced ATP-release [76, 80] that would increase both the area invaded by Ca^2+^ waves and the [Ca^2+^]_C_ increments in individual keratinocytes.

Testing these hypotheses required further pharmacological interventions, as summarized hereafter.

### Effect of thapsigargin

To corroborate the involvement of Ca^2+^ release from the ER downstream of P2Y receptor activation by extracellular ATP, we microinjected 4 µl of VS containing 400 nM of thapsigargin (previously dissolved in DMSO), a potent, cell-permeable, selective and irreversible inhibitor of the sarcoplasmic and endoplasmic reticulum Ca^2+^-ATPase family (SERCA pumps) [81]. As expected, thapsigargin caused a significant increase in the basal level of GCaMP6s fluorescence, *F*_0_, from 190 ± 20 gray levels (GLs) in control conditions to 335 ± 5GLs in thapsigargin (P=0.01, two-sample t-test) due to uncompensated passive leakage of Ca^2+^ from the ER. Thapsigargin caused Ca^2+^ waves to propagate over smaller distances, significantly reducing the equivalent radius *R* (*t*) of the invaded area (**Figure 15 a**). Moreover, the increased *F*_0_ values caused an evident reduction of Δ*F*(*t*)/*F*_0_ bystander responses at any order of distance and consequent significantly reduced *a*_*s*.*s*_. (**Figure 15 d-e**). Also the area *I* under Δ*F*(*t*)/*F*_0_ traces and their slope *S* at the inflection point were significantly reduced in thapsigargin vs. controls at all distances from the photodamage site (**Figure 15 f, g**). Data in **Figure 15** relate to experiments conducted in *n*=3 (control) *n*=2 (thapsigargin) non-overlapping areas of earlobe skin in *m*=1 mouse.

**Figure 15:**
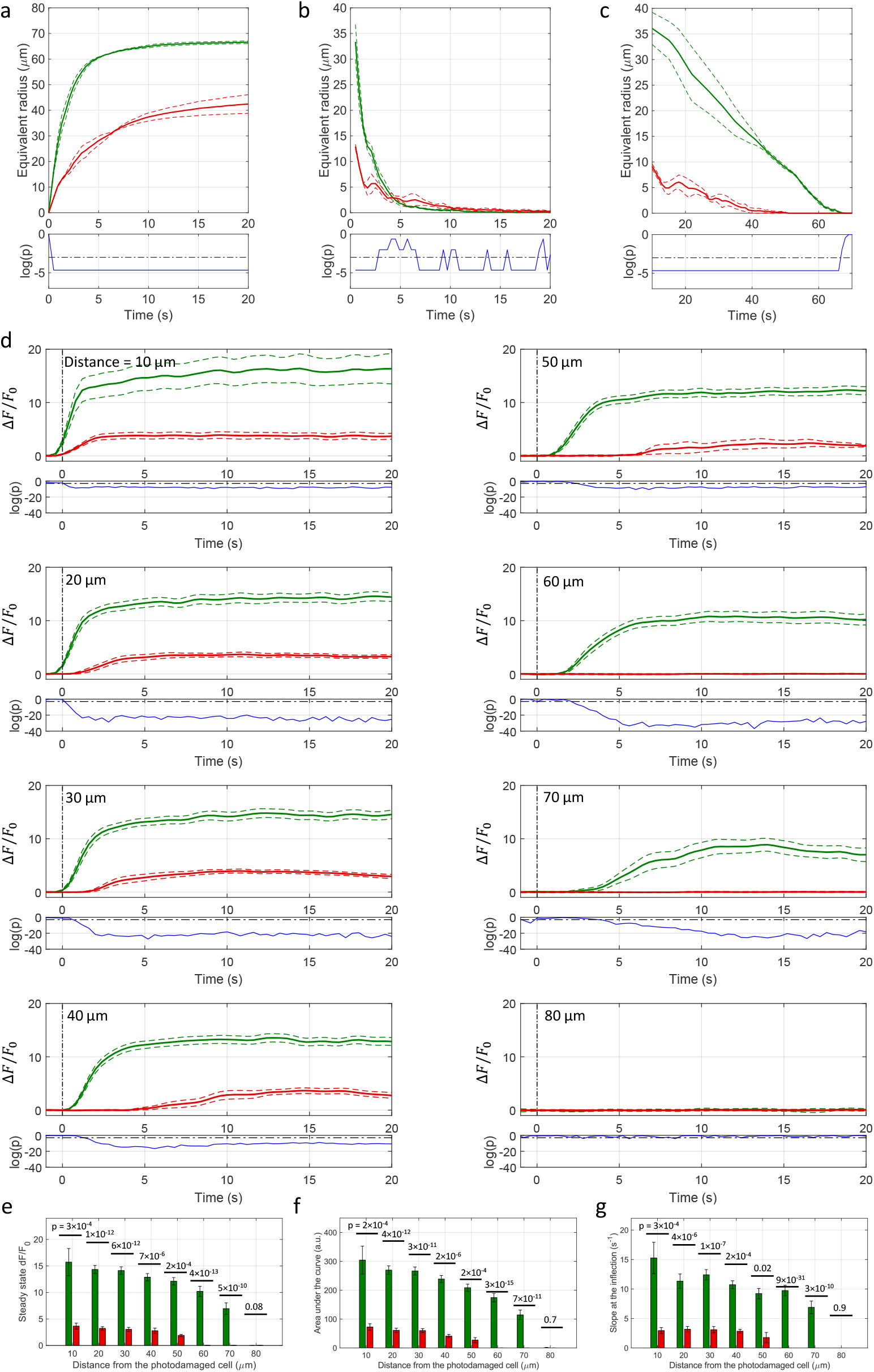
Effect of thapsigargin. (**a**) Equivalent radius of the area invaded by Ca^2+^ waves as a function of time after photodamage; speed of the expanding wave as function of time (**b**) and of the equivalent radius (**c**). (**d**) Δ*F*(*t*)/*F*_0_ responses of bystander keratinocytes at increasing distance from the photodamage site. In each panel, the vertical black dash-dotted line at 0 s marks the end of the 0.5 s photodamage time interval. Data in (**a-d**) are mean (solid line) ± s.e.m. (dashed line) in control conditions (green) and after thapsigargin (440 nM) microinjection (red). Point-by-point p-values (p; Wilcoxon Rank Sum test for **a-c**; two-sample t-test for **d**) are shown on a logarithmic scale below each graph (blue traces); p < 0.05 (horizontal black dash-dotted line) indicates statistical significance. (**e**) Amplitude (*a*_*s*.*s*._) of the Δ*F*(*t*)/*F*_0_ signal at steady state. (**f**) Area (*I*) under the Δ*F*(*t*)/*F*_0_ trace, computed between 0 and 20 s. (**g**) Slope (*S*) of the Δ*F*(*t*)/*F*_0_ trace at the inflection point. Data in (**e-g**) are mean ± s.e.m. vs. bystander cell distance from the photodamage site in control conditions (green) and after thapsigargin microinjection (red). P-values are shown above each pair of bars (two-sample t-test).

### Effect of connexin and pannexin inhibitors

Carbenoxolone (CBX) is a broad-spectrum inhibitor of Panx1 channels [73, 74] that inhibits also connexin-made channel (i.e. HCs and IGJCs) [82]. Recently, CBX has been shown to inhibit also IP_3_-dependent Ca^2+^ release from the ER in endothelial cells of rat mesenteric artery [83]. We injected 4 µl of VS containing CBX (400 µM), which reduced significantly both the area A invaded by Ca^2+^ waves following focal photodamage (**Figure 16 a**) and the Δ*F*(*t*)/*F*_0_ responses in all bystander keratinocytes, except those adjected to the photodamage site (**Figure 16 d**). The parameters a_*s*.*s*._, *I* and S were significantly reduced in CBX compared to controls at *d* > 10 μm, and the arrest distance *d*_stop_ decreased from 80 μm (control) to 70 μm (CBX) (**Figure 16 e-g**). These experiments were conducted in *n*=3 (control) and *n*=4 (CBX) non-overlapping areas of earlobe skin in *m*=1 mouse.

**Figure 16:**
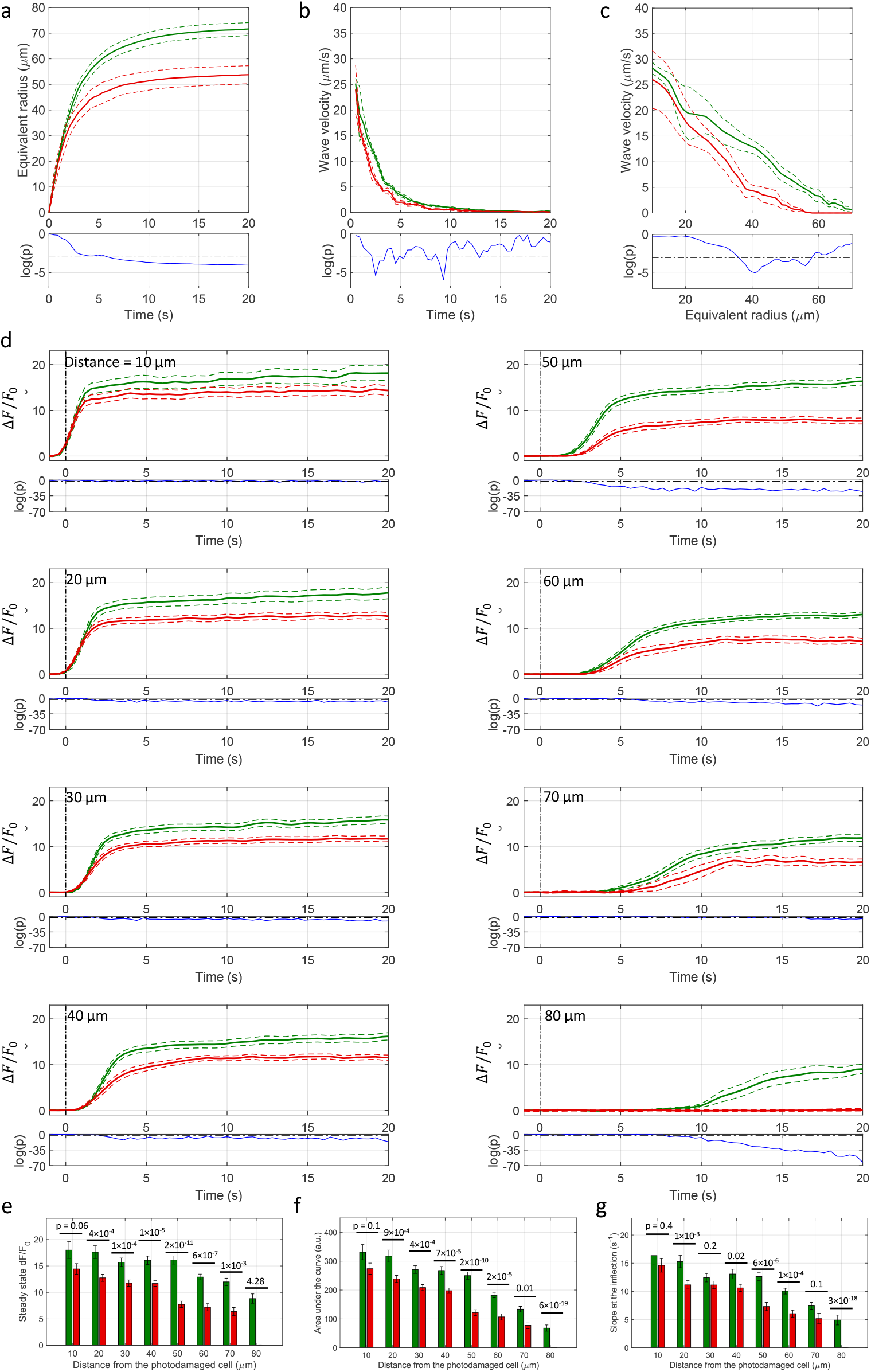
Effect of carbenoxolone (CBX). (**a**) Equivalent radius of the area invaded by Ca^2+^ waves as a function of time after photodamage; speed of the expanding wave as function of time (**b**) and of the equivalent radius (**c**). (**d**) Δ*F*(*t*)/*F*_0_ responses of bystander keratinocytes at increasing distance from the photodamage site. In each panel, the vertical black dash-dotted line at 0 s marks the end of the 0.5 s photodamage time interval. Data in (**a-d**) are mean (solid line) ± s.e.m. (dashed line) in control conditions (green) and after CBX (400 μM) microinjection (red). Point-by-point p-values (p; Wilcoxon Rank Sum test for **a-c**; two-sample t-test for **d**) are shown on a logarithmic scale below each graph (blue traces); p < 0.05 (horizontal black dash-dotted line) indicates statistical significance. (**e**) Amplitude (*a*_*s*.*s*._) of the Δ*F*(*t*)/*F*_0_ signal at steady state. (**f**) Area (*I*) under the Δ*F*(*t*)/*F*_0_ trace, computed between 0 and 20 s. (**g**) Slope (*S*) of the Δ*F*(*t*)/*F*_0_ trace at the inflection point. Data in (**e-g**) are mean ± s.e.m. vs. bystander cell distance from the photodamage site in control conditions (green) and after CBX microinjection (red). P-values are shown above each pair of bars (two-sample t-test).

To discriminate between Panx1 and connexin-made channels, we microinjected 4 µl of VS containing 4 mM of probenecid, which affects primarily Panx1 channels [84]. Probenecid modified slightly the time course of the equivalent radius *R* (*t*) without affecting significantly the area invaded by the Ca^2+^ waves at their maximal expansion (**Figure 17 a**). In addition, Δ*F*(*t*)/*F*_0_ signals were largely unaffected, with the exception of bystander keratinocytes located at 40 μm ≤ *d* ≤ 50 μm where responses appeared increased rather than decreased by the drug (**Figure 17 d**). Only in this range of distances, the parameters a_*s*.*s*._, *I* and S were significantly larger in probenecid compared to controls, and the arrest distance *d*_stop_ was not affected (**Figure 17 e-g**). These experiments were conducted in *n*=3 (control) and n=3 (Probenecid) non-overlapping areas of earlobe skin in *m*=1 mouse and lead us to conclude that (i) Panx1 channels were not involved in the Ca^2+^ signaling triggered by focal photodamage and (ii) the effect of CBX may reflect inhibition of connexin-made channel [82] and/or IP_3_-dependent Ca^2+^ release from the ER [83].

**Figure 17:**
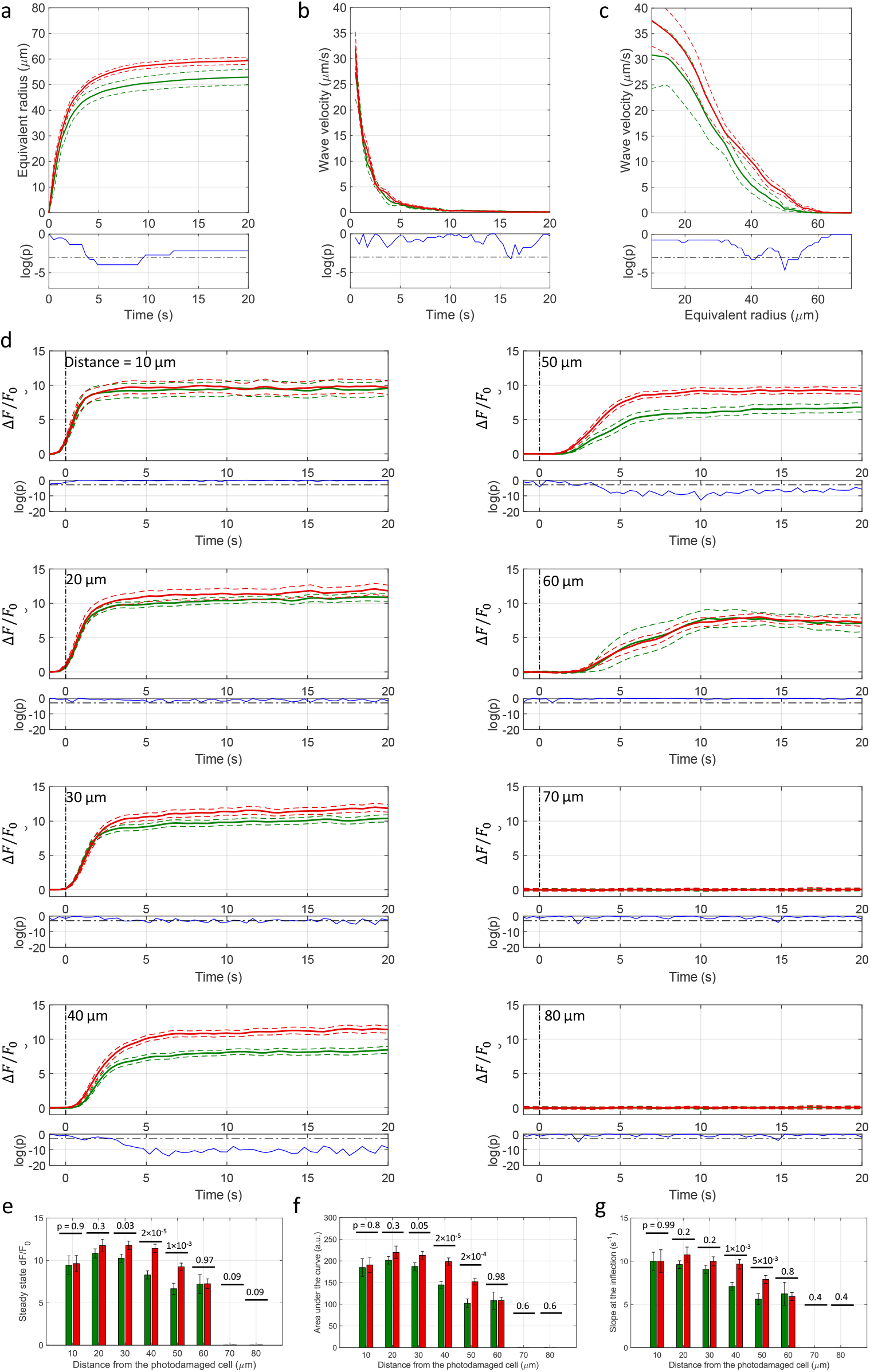
Effect of probenecid. (**a**) Equivalent radius of the area invaded by Ca^2+^ waves as a function of time after photodamage; speed of the expanding wave as function of time (**b**) and of the equivalent radius (**c**). (**d**) Δ*F*(*t*)/*F*_0_ responses of bystander keratinocytes at increasing distance from the photodamage site. In each panel, the vertical black dash-dotted line at 0 s marks the end of the 0.5 s photodamage time interval. Data in (**a-d**) are mean (solid line) ± s.e.m. (dashed line) in control conditions (green) and after probenecid (4 mM) microinjection (red). Point-by-point p-values (p; Wilcoxon Rank Sum test for **a-c**; two-sample t-test for **d**) are shown on a logarithmic scale below each graph (blue traces); p < 0.05 (horizontal black dash-dotted line) indicates statistical significance. (**e**) Amplitude (*a*_*s*.*s*._) of the Δ*F*(*t*)/*F*_0_ signal at steady state. (**f**) Area (*I*) under the Δ*F*(*t*)/*F*_0_ trace, computed between 0 and 20 s. (**g**) Slope (*S*) of the Δ*F*(*t*)/*F*_0_ trace at the inflection point. Data in (**e-g**) are mean ± s.e.m. vs. bystander cell distance from the photodamage site in control conditions (gray) and after probenecid microinjection (light green). P-values are shown above each pair of bars (two-sample t-test).

To discriminate between connexin HCs and IGJCs, we microinjected 4 µl of VS containing 200 µM TAT-Gap19, a peptide that selectively inhibits Cx43 HCs [85]. The area invaded by Ca^2+^ waves did not display statistically significant differences in the presence of TAT-Gap19 compared to controls (**Figure 18 a**), whereas Δ*F*(*t*)/*F*_0_ signals in bystander keratinocytes were attenuated at larger distances (60, 70 µm) (**Figure 18 d**).

**Figure 18:**
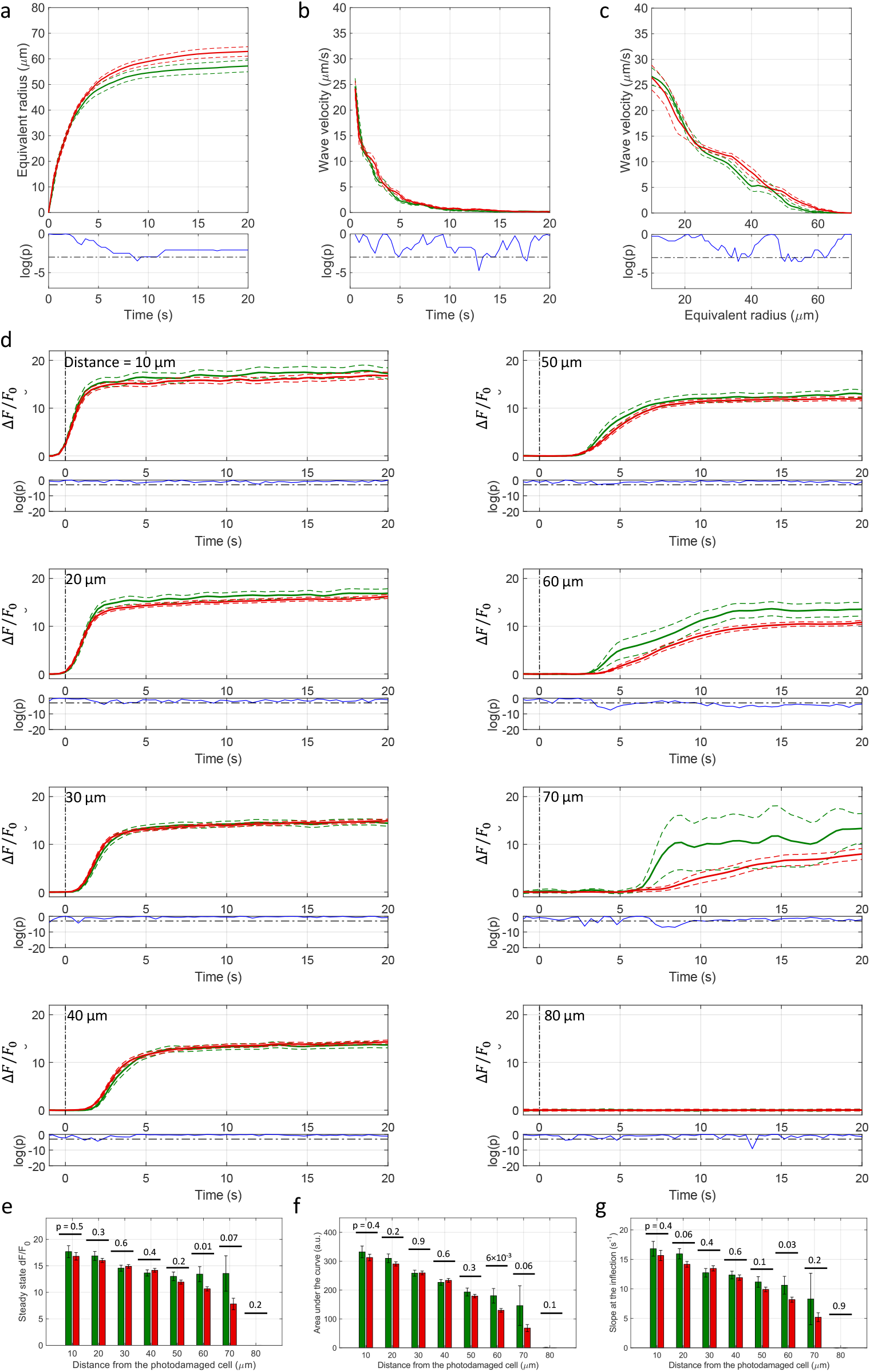
Effect of TAT-Gap19. (**a**) Equivalent radius of the area invaded by Ca^2+^ waves as a function of time after photodamage; speed of the expanding wave as function of time (**b**) and of the equivalent radius (**c**). (**d**) Δ*F*(*t*)/*F*_0_ responses of bystander keratinocytes at increasing distance from the photodamage site. In each panel, the vertical black dash-dotted line at 0 s marks the end of the 0.5 s photodamage time interval. Data in (**a-d**) are mean (solid line) ± s.e.m. (dashed line) in control conditions (green) and after TAT-gap19 (200 μM) microinjection (red). Point-by-point p-values (p; Wilcoxon Rank Sum test for **a-c**; two-sample t-test for **d**) are shown on a logarithmic scale below each graph (blue traces); p < 0.05 (horizontal black dash-dotted line) indicates statistical significance. (**e**) Amplitude (*a*_*s*.*s*._) of the Δ*F*(*t*)/*F*_0_ signal at steady state. (**f**) Area (*I*) under the Δ*F*(*t*)/*F*_0_ trace, computed between 0 and 20 s. (**g**) Slope (*S*) of the Δ*F*(*t*)/*F*_0_ trace at the inflection point. Data in (**e-g**) are mean ± s.e.m. vs. bystander cell distance from the photodamage site in control conditions (green) and after TAT-gap19 microinjection (red). P-values are shown above each pair of bars (two-sample t-test).

The parameters a_*s*.*s*._, *I* and S were significantly reduced in Tat-Gap19 compared to controls at *d* > 50 μm, but the arrest distance *d*_stop_ was not affected (**Figure 18 e-g**). These experiments were conducted in *n*=3 (control) *n*=6 (TAT-Gap19) non-overlapping areas of earlobe skin in *m*=1 mouse.

### Summary of experimental results

The major effects of drugs that interfere with degradation of extracellular ATP or P2 purinoceptors suggest that Ca^2+^ waves in the photodamaged epidermis are primarily due to release of ATP from the target cell, whose plasma membrane integrity was compromised by laser irradiation. The ineffectiveness of probenecid suggests pannexin channels have no role. As Δ*F*(*t*)/*F*_0_ signals in bystander keratinocytes were augmented by exposure to the Ca^2+^ chelator EGTA in the extracellular medium, the corresponding transient increments of the [Ca^2+^]_c_ should be ascribed primarily to Ca^2+^ release from the ER, downstream of ATP binding to P2Y purinoceptors (P2Y_1_, P2Y_2_, P2Y_4_ and P2Y_6_), with Ca^2+^ entry through plasma membrane channels playing a comparatively negligible role. The effect of thapsigargin (a well-known inhibitor of SERCA pumps) supports this conclusion. The limited effect of TAT-Gap19 suggest ATP-dependent ATP release though connexin HCs played a minor role, affecting Ca^2+^ wave propagation only at larger distances, where the concentration of ATP released from the photodamaged cell was reduced by the combined effect of passive diffusion and hydrolysis due to the action of ecto-ATPases. The fact that CBX exerted a stronger inhibitory action compared to TAT-Gap19 suggests IGJCs played a more prevalent role than Cx43 HCs in Ca^2+^ wave propagation. However, CBX has been recently reported to interference also with IP_3_ receptors [86], which would instead lend further support to a fundamental role played by the P2Y-dependent pathway and IP_3_-dependent Ca^2+^ release from intracellular stores.

## DISCUSSION

We have shown here that ATP is released from epidermal keratinocytes of live anesthetized mice following intradermal photodamage caused by intense pulsed IR laser radiation. Kumamoto *et al*. obtained results compatible with those reported here using point laser stimulation of keratinocytes in *ex vivo* human epidermis. In particular, they observed different responses in the various layers of the epidermis after stimulating a keratinocyte of the *stratum granulosum*, with higher responses in the basal layer [28].

### Significance of these findings for would healing and skin therapy

It is tempting to speculate that response coordination after injury or laser treatment occurs via propagation of the ATP-dependent intercellular Ca^2+^ waves we visualized and analyzed in this article. ATP is well known as one of the damage-associated molecular patterns (DAMPs) that, once released from injured cells, activate the immune system [87]. In the epidermis, Langerhans cells, characterized by the expression of Langerin (CD207) and CD1a, are considered to be macrophages that retain the function of dendritic cells, playing a clear role of antigen-presenting cells in skin inflammation, triggering a series of immune responses by migrating from the epidermis to the lymph nodes and presenting antigens to T-regulatory cells [88]. A Langerhans cell line (XS106) was shown to express mRNA for P2X_1_, P2X_7_, P2Y_1_, P2Y_2_, P2Y_4_, and P2Y_11_ receptors [89], which is consistent with the increase in GCaMP6s fluorescence in Langerhans cells we observed in some of the experiments. P2Y receptors regulate many aspects of immune cell function, including phagocytosis and killing of pathogens, antigen presentation, chemotaxis, degranulation, cytokine production, and lymphocyte activation [90]. Therefore, it would be interesting, in future, studies, to trace the movements of the Langerhans cells invested by damage-evoked Ca^2+^ waves.

Besides alerting the body about danger and stimulating the immune system in order to initiate the immune response, DAMPs also promote the regeneration process [87]. Our results clearly indicate that P2YR were activated by ATP downstream of photodamage. P2YR regulate cell proliferation, differentiation and migration [91-93] and, are also involved in wound healing [24]. A prime target of the associated signaling activity is likely to be the extracellular matrix (ECM). In cultured keratinocytes, ATP caused a rapid and strong but transient activation of hyaluronan synthase 2 (*HAS2*) expression (via PKC-, CaMKII-, MAPK- and CREB-dependent pathways) by activating P2Y_2_ receptors [94]. Hyaluronan, also named hyaluronic acid, is a major component of the ECM, and its synthesis is rapidly upregulated after tissue wounding [95]. Thus, epidermal insults are associated with extracellular ATP release, as well as hyaluronan synthesis, and the two phenomena are linked [94].

The micro-regions targeted by high intensity laser-induced intradermal photodamage are thought to promote formation of new dermal collagen and elastin during the repair process [32-35]. Fibroblasts are well known not only for being the major producers of extracellular matrix, but also a cell type actively involved in synthesis of inflammatory mediators and tissue repair. ATP or UTP increased the levels of ECM-related proteins through the activation of P2Y2R in mouse fibroblasts [96]. In our experimental model, Ca^2+^ waves reached into the fibroblast-populated collagen matrix, and this event may activate and coordinate the keratinocyte-fibroblast interaction that is a key to wound healing and skin remodeling [97, 98].

We did not analyze in detail the wining phase of Ca^2+^ waves that, in our experiments, lasted up to 120 minutes, with a reduction of half the maximum invaded area after ∼20 minutes from the photodamage. This latter part of the Ca^2+^ dynamics may involve a variety of other processes we did not take into account. Indeed, numerous other molecular players contribute to epidermal Ca^2+^ homeostats, including Ca^2+^-sensing receptor, transient receptor potential channels, store-operated calcium entry channel Orai1, endoplasmic Ca^2+^ depletion sensor (stromal interaction molecule 1 [STIM1]), and voltage-gated calcium channels such as L-type calcium channels [10, 99]. In addition, further experiments are required to better characterize the dynamics of ATP after it is released into the extracellular space and is degraded rapidly to ADP, AMP, and further to adenosine [60], which (with the exception of AMP) activate specific receptors even in a self-sustaining fashion [100].

## Author contributions

*Conceptualization:* F.M.; *Data Curation:* V.D.; *Funding Acquisition:* FM. *Investigation:* V.D., C.P.; *Methodology:* C.N., M.R., F.S., M.B.; *Project Administration:* F.M.; *Resources:* M.R.; F.S.; *Software:* V.D., M.B., C.D.C., D.A.C.; *Supervision:* F.M., D.A.C., M.B.; *Validation:* F.M.; *Visualization:* V.D., C.P.; *Writing - Original Draft Preparation:* V.D., C.P., F.M.; *Writing - Review and Editing:* F.M..

